# Leveraging whole-genome re-sequencing for diversity, population structure, and a public mid-density genotyping enrichment panel in crimson clover (*Trifolium incarnatum L*.) for breeding purposes

**DOI:** 10.64898/2026.01.26.701603

**Authors:** Mark Philip Castillo, Oluwaseye Gideon Oyebode, Jayson Talag, Victoria Bunting, Navneet Kaur, Paul Doran, Kerrie Barry, Jeremy Schmutz, Brandon Schlautman, Alan Humphries, Kioumars Ghamkhar, Virginia Moore, Esteban Rios, Alex Harkess, Marnin Wolfe

## Abstract

Crimson clover *(Trifolium incarnatum L.)* is an obligately outcrossing, cool-season annual legume valued for forage and cover cropping, yet genomic resources to support systematic improvement are limited. We performed the first and most comprehensive whole-genome resequencing (WGR) of global crimson clover germplasm to (i) characterize diversity and population structure and (ii) develop a public mid-density enrichment capture panel for breeding applications. A core set of 45 accessions sequenced at ∼50X generated 5.84 million variants, while 149 additional accessions sequenced at ∼2.54X yielded 17.05 million variants. After stringent filtering, we retained 542,790 high-confidence SNPs from the high-coverage dataset and ∼2.4 million from the low-pass cohort. Population analyses (PCA, ADMIXTURE) revealed compact clustering of cultivars, broader dispersion of wild and uncertain-status accessions, and low overall differentiation (FST = 0.0105) with excess heterozygosity (FIS = -0.0592), consistent with obligate outcrossing. Guided by these resources, we designed a 28,913-SNP TWIST hybrid-capture panel enriched for genic regions and evenly distributed across seven chromosomes. This panel is being deployed within Auburn University’s crimson clover breeding program to support population improvement and cultivar development. The resulting genomic resources provide a reproducible, mid-density genotyping platform for trait discovery, predictive breeding, and diversity monitoring. Together, these advances bring crimson clover genomic resources on par with other legumes such as soybean (*Glycine max (L.) Merr*.) and alfalfa (*Medicago sativa*), establishing a robust foundation for genomics-assisted improvement of this key cover and forage crop in U.S. sustainable agriculture.

**CORE IDEAS:**

- Whole-genome re-sequencing of 194 crimson clover accessions revealed >21 M variants.
- High-confidence SNP catalogs from 50X and 2X data enable cost-effective genotyping.
- Genetic diversity is weakly structured, with cultivars clustering narrowly by origin.
- A 28,913 SNP enrichment panel delivers uniform genome coverage and >75% genic content.
- These genomic tools accelerate GWAS, genomic selection, and breeding innovation.

## Introduction

Crimson clover *(Trifolium incarnatum L.)* is a cool-season, obligately outcrossing legume valued worldwide as both a forage and cover crop due to its high biomass production, strong nitrogen fixation capacity, and contributions to soil fertility and ecosystem resilience (Barnes et al, 1995; Moore et al., 2020). By reducing reliance on synthetic fertilizers and enhancing nutrient cycling, crimson clover plays a vital role in sustainable intensification across diverse cropping systems. Recent surveys underscore its growing agricultural importance, such as the 2022-2023 National Cover Crop Survey, which ranks crimson clover first among legumes and fourth among all cover-crop species planted in U.S. fields, behind only cereal rye (*Secale cereale L*.), radish (*Raphanus sativus L*.), and winter wheat (*Triticum aestivum L*.), and with many growers reporting more than a decade of continuous use, reflecting both agronomic reliability and long term soil benefits (CTIC,SARE,&ASTA, 2023). Today, crimson clover seed production is approximately 2.6-4.4 million kg of seed annually, valued at $3.9–$6.2 million in farm-gate sales. Production is concentrated in Oregon’s Willamette Valley, which now supplies nearly all U.S. crimson clover seed. Between 2018 and 2020, harvested area in Oregon ranged from 2,315 to 3,650 ha, with average yields of approximately 1,120 kg ha−^1^ (Oregon State University Extension Service, 2021).

Despite its agronomic and ecological importance, crimson clover has historically lagged far behind other forage and grain legumes in genomic resource development. While model and crop legumes such as soybean, alfalfa, and red clover (*Trifolium pratense*) now benefit from chromosome-scale reference genomes, dense SNP arrays, and large-scale diversity studies that have transformed molecular breeding (Bickhart et al., 2021; Varshney et al., 2021), comparable genomic resources for crimson clover remain limited. Within the genus *Trifolium*, most genomic studies have focused on red and white clover (*Trifolium repens*). Red clover now possesses multiple chromosome-level assemblies supporting GWAS, QTL mapping, and genomic selection for key forage agronomic traits (De Vega et al., 2015; Yan et al., 2022; Bickhart et al., 2022). While white clover, an allotetraploid, has similarly high-contiguity genome assemblies that reveal recombination landscapes and inform polyploid prediction (Kuo et al.,2024). In contrast, our only knownledge about crimson clover includes a C-value genome size estimate (∼652 Mb; 2n = 14) and an early AFLP-based diversity surveys (Steiner et al., 1998; Vižintin et al., 2006; Ellison et al., 2006). The absence of whole genome assemblies and genome-wide markers is constraining breeding progress in crimson clover.

Reduced-representation methods such as SSRs or genotyping-by-sequencing (GBS) have provided only partial insight, capturing only a small fraction of the genome and producing marker sets that are ofter inconsistent across studies. Because GBS relies on restriction enzyme digestion, it often suffers from missing data, locus dropout, and batch effects that limiting reproducibility and downstream application (Elshire et al., 2011; Poland & Rife, 2012). In contrast, whole-genome resequencing (WGR) offers comprehensive variant discovery across coding and noncoding regions, enabling dense SNP catalogs, accurate estimates of linkage disequilibrium, and the foundation for genomic tools that can accelerate breeding (Tester & Langridge, 2010; Varshney et al., 2014). Expanding such genomic resources for crimson clover is therefore a critical need.

While WGR remains too costly for routine breeding applications, it provides the discovery layer for designing efficient, reduced-representation genotyping tools. One such approach is hybrid-capture enrichment panels, probe-based genotyping platforms that selectively sequence thousands of predefined genomic regions. Hybrid capture employs short synthetic oligonucleotide “baits” that hybridize to target loci, enabling reproducible, genome-wide SNP recovery at moderate cost (Lemmon et al., 2012; Twist Bioscience, 2023). Compared with GBS, enrichment panels offer consistent marker sets, reduced missing data, and improved cross-study comparability, making them ideally suited for genomic selection, GWAS, and germplasm characterization.

To fill this gap, we conducted the first WGR study of crimson clover germplasm. We applied a two-tiered re-sequencing strategy with a core set sequenced at high (∼50X) coverage and a broader set at low (∼2X) coverage; this approach balances coverage, resolution and cost. The specific objectives were to: (i) generate a comprehensive SNP resource to support genomic tool development; (ii) characterize genetic diversity, population structure, and private variation across a global panel; and (iii) develop a publicly available, mid-density genome-wide hybrid-capture enrichment panel (∼28K SNPs) optimized for crimson clover breeding applications. Together, these efforts establish the first genome-wide variant resource for crimson clover and provides a bridge between discovery genomics and applied breeding, bringing this agronomically important legume into the genomics era.

## Materials and Methods

### Plant Material and Sample Collection

A total of 45 distinct crimson clover accessions were collected for this study, representing a mixture of synthetic populations and maternal half-sib families. Specifically, 40 accessions were obtained from the USDA National Plant Germplasm System (NPGS), encompassing plant introductions (PIs) from diverse geographic origins (Figure 3). NPGS accessions generally represent small synthetic populations of open-pollinated individuals with unknown effective population sizes, reflecting natural diversity. In contrast, accessions provided by Auburn University (AU) (2 accessions) and the University of Florida (UF) (2 accessions) were derived from maternal half-sib families, each representing progeny of open-pollinated plants selected within breeding nurseries. The AU accessions consisted of one polycross family (AUCC14) originating from a cross between the cultivars AU Robin and AU Reseeding. The UF accessions included two half-sib families (UFCC17 and UFEarlyWhite). In addition, the commercial cultivar Dixie (1 accession) was included as a replicated check. Seeds were sown in October 2022, with 12 seeds per family in 72-cell trays. After six weeks, seedlings were transplanted to the E.V. Smith Research Station – Plant Breeding Unit in Tallassee, Alabama, USA (32.5117° N, 85.8925° W). Plants were spaced 18 inches apart in a nursery layout following an augmented design, with the cultivar ‘Dixie’ included as a check in each row, totaling 350 plants at establishment. Following a winter freeze event, 194 plants survived and were retained for tissue sampling, consisting of 154 PI plants, 18 AU plants, 5 UF plants, and 17 ‘Dixie’ check plants. Young leaf tissue was collected in early 2023 during late vegetative and early reproductive stages, flash-frozen in liquid nitrogen, and stored at –80 °C until DNA extraction.

### DNA Extraction

Genomic DNA was extracted from lyophilized tissues using a modified cetyltrimethylammonium bromide (CTAB) protocol optimized for leguminous species (Doyle 1990). This protocol incorporated additional steps to remove polysaccharides and secondary metabolites. DNA quantity and quality were assessed using a NanoDrop 2000 spectrophotometer (Thermo Fisher Scientific) and Qubit 4 Fluorometer (Invitrogen), following standard manufacturer protocols.

### Two-tiered sequencing strategy

The 194 individual plants (unique genotypes) were sequenced as part of a two-tiered whole-genome resequencing (WGR) strategy. For each of the 45 accessions, up to ten plants were transplanted to the field to capture within-accession diversity. Because crimson clover is genetically heterogeneous, multiple individuals per accession were evaluated to represent population variation. Three phenotypic traits were scored: plant height, vigor (visual score, 3 > 5 > 7), and SPAD (a proxy for chlorophyll and nitrogen content). A linear mixed model was fitted for each trait to account for field blocking effects, and best linear unbiased predictors (BLUPs) were obtained. A Smith-Hazel selection index (Smith, 1936; Hazel, 1943) was constructed to combine these traits into a single composite score. The index used economic weights of 0.5,0.3, and 0.2 for height, vigor, and SPAD, respectively, emphasizing biomass and plant robustness. Because pedigree or genomic relationships were not yet available, the index was applied at the phenotypic level to identify the best individuals within each family. For each accession or family, one top-ranked plant based on the index was selected for high-coverage sequencing (∼50X), while the remaining individuals were sequenced at low coverage (∼2X) to maximize population representation. In total, 45 plants were sequenced at high coverage (50X) and 149 at low coverage (2X), forming a discovery-plus-diversity dataset.

### Library preparation and sequencing

Genomic DNA from all samples was randomly sheared to ∼500 bp and used to prepare Illumina paired-end libraries. Sequencing was performed on the Illumina NovaSeq 6000 and Illumina NovaSeq X, generating 150 bp paired-end reads. Quality control and adapter trimming were applied to raw reads prior to alignment to the *Trifolium pratense* reference genome (GCF_020283565.1_ARS_RC_1.1) using BWA-MEM 07.17. Variant calling was conducted using SAMtools 1.11 and mpileup. All data, including FASTQs, BAMs, and VCFs, are to be publicly accessible via the Department of Energy (DOE) Joint Genome Institure (JGI) Genome Portal under Proposal ID 509088.

### Sequence Quality Control, Read Alignment, and Variant Discovery

Raw Illumina NovaSeq 6000 and Illumina NovaSeq X 2×151 bp reads were initially processed using DOE JGI following *Resequencing SOP 1099.3* (JGI, 2023). Read filtering and quality improvement were performed using the BBTools v39.03 RQCFilter2 workflow (Bushnell, 2014). This pipeline includes optical and PCR duplicate removal (Clumpify), adapter trimming, removal of terminal poly-G sequencing artifacts, right-end quality trimming below Q6, and filtering of reads containing ambiguous bases or failing minimum-length requirements. The workflow also screens against a comprehensive contamination database (PhiX, microbial, organellar, and animal references), though no contaminant reads were detected across libraries. After filtering, the retained high-quality reads represented the non-redundant, informative sequence content of each library. High-quality reads were aligned to the *Trifolium pratense* reference genome (GCF_020283565.1 ARS_RC_1.1; Bickhart et al., 2022) using BWA-MEM v0.7.17 with default settings (Li & Durbin, 2009). Alignment sorting, indexing, and processing were performed using SAMtools v1.11 (Li et al., 2009) and variant calling was conducted on the Auburn University High-Performance Computing (AU-HPC) system using bcftools 1.11 mpileup followed by bcftools call to generate per-sample variant call files (VCFs). Individual VCFs were subsequently merged through joint variant calling to produce a unified multi-sample dataset for each coverage level. Variants were filtered with bcftools filter using quality and depth thresholds (QUAL > 30 & DP > 10), retaining only biallelic SNPs. Additional filters included a minor allele frequency (MAF) ≥ 0.05 and a missing genotype rate ≤ 0.10, ensuring that retained sites were genotyped in at least 90 % of individuals. The high-coverage (50X) and low-coverage (2X) datasets were processed independently through similar variant-discovery and filtering pipelines implemented with bcftools 1.18 and PLINK 1.9. After variant normalization with bcftools norm, haplotype phasing and genotype imputation were performed using Beagle 5.4 (Browning & Browning, 2016). Complete filtering parameters and workflow details are provided in Tables 2.1 and 2.2.

**Table 1.** SNP filtering strategy for the 50X WGR dataset of 45 *T. incarnatum* accessions, yielding 542,790 high-confidence SNPs for downstream analyses.

| Filtering Step | Criteria Applied | Remaining SNPs | Percentage (%) |
| --- | --- | --- | --- |
| Total Variants Identified | Initial variant calling (50X coverage) | 5,843,860 | 100 |
| SNPs Identified | SNP subset from total variants | 4,958,826 | 84.86 |
| 30% Missing Data Filter Applied | Removal of SNPs with >30% missing data | 1698625 | 29.07 |
| Minor Allele Frequency (MAF < 0.05) | MAF threshold set at 0.05 | 542790 | 9.29 |
| Beagle 5.4 Imputation Applied | Imputation for missing genotypes | 542790 | 9.29 |
| Final High-Confidence SNPs | Quality-controlled dataset | 542790 | 9.29 |

**Table 2.** Chromosomal distribution and SNP density in crimson clover based on both high-confidence and low confidence SNPs.

| Chromosome | Size(Mb)(a) | SNP Number |  |  |  |
| --- | --- | --- | --- | --- | --- |
|  |  | ~50X Coverage |  | ~2X Coverage |  |
|  |  | SNP number | SNP density per Mb | SNP number | SNP density per Mb |
| 1 | 53.59 | 84,382 | 1574.6 | 353,540 | 6597.3 |
| 2 | 74.48 | 93,614 | 1256.9 | 402,557 | 5404.8 |
| 3 | 54.40 | 90,252 | 1659.1 | 389,206 | 7154.6 |
| 4 | 60.49 | 77,948 | 1288.5 | 370,493 | 6124.2 |
| 5 | 57.90 | 55,178 | 952.9 | 268,064 | 4629.2 |
| 6 | 46.12 | 64,766 | 1404.4 | 286,864 | 6220.6 |
| 7 | 59.21 | 76,650 | 1294.5 | 340,744 | 5754.7 |
| Average | 58.03 | 77,541.40 | 1347.27 | 2,411,468 | 41885.4 |

<sup>a</sup>Based on the reference genome of red clover Genome assembly ARS\_RC\_1.1, Bickhart et al. 2022

### Targeted Enrichment Panel Design

Workflows were anchored to the red clover (ARS_RC_1.1) reference assembly (RefSeq GCF_020283565.1). Coordinate conventions were harmonized across file formats by treating VCF positions as 1-based and BED intervals as 0-based end-exclusive, and by enforcing exact chromosome naming consistency with the FASTA index (NC_060059.1–NC_060065.1).

Gene models from the ARS_RC_1.1 GFF annotation were merged to define a comprehensive “genic union,” which included entire gene bodies and their associated exon, CDS, and UTR features. Candidate SNP discovery integrated three independent variant sources: a high-coverage (∼50X) JGI panel, a low-coverage (∼2X) JGI panel, and a panel of 6,803 genome-wide SNPs identified in a separate genotyping project in our breeding program in 2023, utilizing Khufu (Korani et al 2021) (https://www.hudsonalpha.org/khufudata/) genotyping in our panel of crimson clover breeding germplasm in 2023. The dataset originated from an intermediate file produced by containing approximately 6,803 marker positions filtered to retain only sites with ≤80% missing data, read depths between 25 and 500, and allele frequencies ranging from 0.05 to 0.95. All VCF files were normalized against the same reference FASTA to ensure consistency. This process involved left-aligning and decomposing multiallelic sites, standardizing REF/ALT allele representations, and then compressing and indexing the outputs prior to joint processing. All analyses were conducted on the Auburn University High Performance Computing (AUHPC). The pipeline utilized version-controlled AUHPC modules, including bcftools/1.20, htslib/1.20, samtools/1.20, bow/2.31.1 (Quinlan & Hall, 2010), and plink2/2.00-alpha (build a5.x or later). Software versions are reported as loaded on AUHPC between July and August 2025 to ensure computational reproducibility and transparency.

Each sequencing strategy produced distinct dataset characteristics and post-NGS specifications requiring customized workflow optimization. In this phase (Figure 12), variant calls from the high-coverage (50X) and low-coverage (2X) JGI datasets were anchored to the red clover reference genome (RefSeq GCF_020283565.1). The two variant call sets were merged with *bcftools merge* to generate a comprehensive variant catalog for probe design. Site-level quality control was implemented with bcftools view and vcftools 0.1.16, retaining only biallelic SNPs with QUAL ≥ 30, minor allele frequency (MAF) ≥ 0.03, and per-site missingness ≤ 0.10. The slightly relaxed MAF threshold followed predictive-breeding panel-design principles to include moderately rare alleles that enhance marker diversity and genome coverage. Strand-ambiguous polymorphisms (A/T and C/G) were identified and excluded using a custom awk/grep filter, while linkage disequilibrium (LD) pruning was carried out in plink 1.9 with a 50-kb window, 5-kb step, and r^2^ ≤ 0.2 (validated at r^2^ ≤ 0.1) to ensure independence among retained variants.

Each variant was annotated as genic or non-genic via bedtools intersect, comparing variant coordinates against a merged genic union GFF3 annotation derived from the ARS_RC_1.1 reference. Overlapping records were resolved deterministically using a short Python 3.10 script that prioritized genic status and higher MAF values. Capture targets were then defined by expanding each site ± 60 bp with bedtools slop, merging overlaps with bedtools merge, and converting outputs into BED3 format. To enforce chromosome-proportional coverage, a bin-center heuristic implemented in Python (pandas + numpy) partitioned each chromosome into equal-length bins and selected the variant nearest the bin midpoint, redistributing gaps iteratively to maintain balanced genomic coverage. The optimized JGI-derived set was unified with a curated Khufu-filtered variant pool using bcftools isec and bedtools merge to collapse overlapping or adjacent intervals. A final “top-up” procedure scripted in Python favored genic features and ensured uniform spacing across the genome. Chromosome-level distribution plots were generated on AUHPC using Python 3.10, CairoSVG ≥ 2.7, and Pillow ≥ 10 within a minimal virtual environment. Visualization scripts parsed FASTA indices and BED3 coordinates to produce high-resolution PNGs labeled with both “Chr01–Chr07” and reference accession identifiers.

In-silico probe validation was conducted using Bowtie2 v2.4.2 (Langmead & Salzberg, 2012). Each ± 60 bp window was converted to a 121-bp synthetic probe sequence with bedtools getfasta, and the reference genome was indexed via bowtie2-build and synchronized with samtools faidx. End-to-end alignments were executed under the --very-sensitive preset with one seed mismatch (-N 1) and a reporting limit of two placements per probe (-k 2). A secondary stress test using local alignments (--very-sensitive-local, -L 20, -N 1, -k 5) identified potential partial or clipped matches. Probes mapping uniquely under both alignment modes were retained for synthesis. Following in-silico validation, the final probe manifest was submitted to TWIST Bioscience (https://www.twistbioscience.com/products/ngs/custom-panels.), where internal quality control confirmed probe balance, hybridization potential, and manufacturing feasibility. TWIST’s verification ensures compatibility with capture chemistry and production, providing an additional layer of validation beyond computational assessment.

### TWIST hybrid capture enrichment panel verification using imputed HapMap genotypes

The TWIST hybrid-capture enrichment panel combined with Khufu low-pass whole-genome sequencing was implemented in the 2024 and 2025 crimson clover breeding germplasm panel of maternal half-sib families conducted across field locations at AU and the UF (genotyping scheme shown at Figure 1), providing a breeding-relevant evaluation of panel performance under applied selection environments. HapMap genotype datasets were generated as part of this workflow. One dataset was derived exclusively from enriched sequencing data, in which variant calling was restricted to SNPs overlapping the TWIST probe intervals defined in the panel BED files. For the enriched-only dataset, the file KHU169_enriched_SNPcalls.hapmap contained 72,181 SNPs genotyped 1,039 lines, with the corresponding imputed dataset provided as KHU169_enriched_SNPcalls.Ihapmap. Genotyping performance was evaluated using this imputed enriched-only HapMap, which represents the final, user-facing genotype matrix produced from Khufu low-pass sequencing (∼3X), targeted capture, and reference-guided imputation. HapMap files were imported into R (data.table), and genotype strings were converted to additive allele dosages (0, 1, or 2 copies of the alternate allele) using marker-specific allele definitions provided in the HapMap alleles field. Non-informative or invalid genotype entries (e.g., missing calls, ambiguous bases, malformed strings, or alleles inconsistent with expected REF/ALT states) were set to NA prior to transformation. Dosage conversion was parallelized across samples (parallel::mclapply) to construct a SNP x sample dosage matrix. Panel completeness was quantified as (i) per-sample call rate, defined as one minus the proportion of missing genotypes across loci, and (ii) per-locus call rate, defined as one minus the proportion of missing genotypes across samples. Minor allele frequency (MAF) was calculated from dosage values as *MAF* = min(*p*, 1 − *p*), *where* 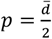 *and* 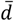 is the mean dosage across samples after excluding missing values Genotyping performance was visualized using histograms and empirical cumulative distribution functions (ECDFs) of call rate, as well as MAF distributions for loci meeting standard usability thresholds (locus call rate ≥ 0.95; MAF ≥ 0.05), implemented using ggplot2.

**Figure 1.**
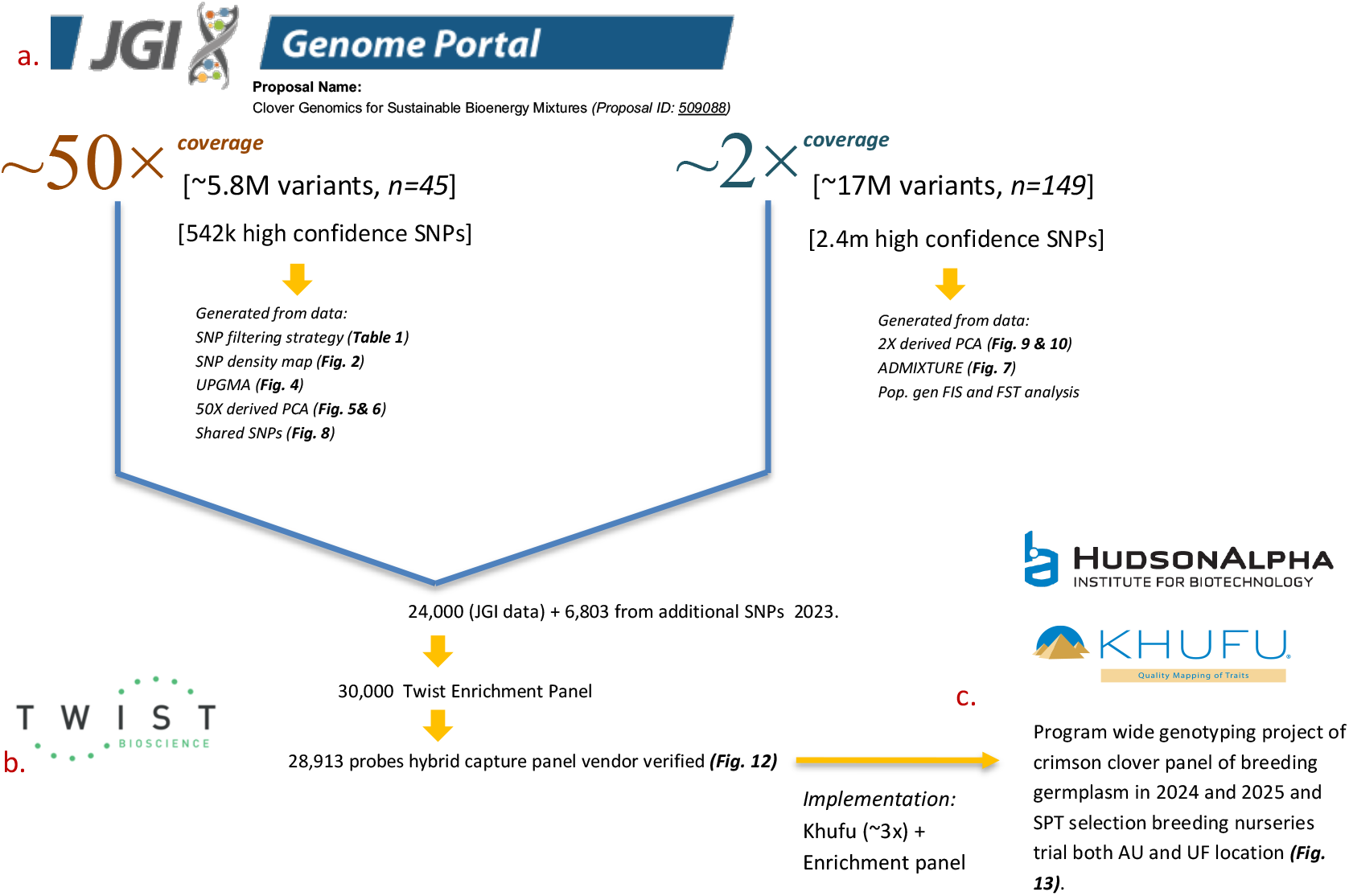
Overview of Auburn crimson clover breeding program genomics workflow and resources. a. JGI NGS service and data storage, b. TWIST DNA synthesis vendor, c. Application of genomic data within the breeding pipeline to support marker discovery.

### Exploratory high coverage (50X) guided imputation to low coverage (2X)

This analysis represents an exploratory approach designed to evaluate how high-coverage and low-coverage datasets can be jointly utilized for downstream genomic studies. It was conducted independently of the enrichment panel and population structure analyses described above but demonstrates a framework for utilizing variant calls across sequencing depths. Variant representations between the high-coverage panel (50X) and the low-coverage cohort (2X) were harmonized against the red clover reference genome (*Trifolium pratense* ARS_RC_1.1; RefSeq GCF_020283565.1). VCFs from both cohorts were left-aligned and normalized using bcftools v1.18, and REF alleles were reconciled to the reference FASTA using the bcftools + fixref plugin. Sites with irreconcilable REF/ALT mismatches were discarded to prevent strand or allele orientation errors. After harmonization, the founder panel comprised 542,790 SNPs with 100% REF matches (no flips, swaps, or unresolved sites), while the 2X target dataset contained 8,205,365 SNPs prior to cross-cohort alignment. To enable guided imputation and ensure allele consistency, the 2X dataset was restricted to sites identical to the founder panel (chromosome, position, REF, ALT). Genotype imputation was performed using Beagle v5.4 (build 22Jul22.46e), with the harmonized 50X founders as the reference haplotype panel and the intersected 2X dataset as the target.

Beagle generated genotype dosages and corresponding dosage R^2^ (DR^2^) accuracy estimates for each SNP. Post-imputation filtering retained variants with DR^2^ ≥ 0.8 and MAF ≥ 0.01 for downstream population analyses. Where permitted by statistical models, continuous dosage values were used instead of hard genotype calls. The final imputed dataset showed near-complete concordance with the founder reference panel, negligible missingness, and consistent allele encodings across samples, supporting the robustness of this guided-imputation strategy. Although this approach necessarily restricts analyses to sites shared between the founder and low-coverage cohorts, thereby underrepresenting rare or private alleles unique to the 2X dataset, it maximizes allele consistency and enables reliable cross-cohort comparisons. This trade-off provides a scalable and cost-effective framework for future *T. incarnatum* sequencing efforts at variable depths.

Imputation quality was assessed using Beagle’s DR^2^ metric. Site-level DR^2^ and allele frequencies (AF) were extracted using bcftools +fill-tags, and the proportion of variants exceeding key accuracy thresholds (DR^2^ ≥ 0.8 and DR^2^ ≥ 0.9) was quantified using standard Unix/AWK pipelines. Visualization was performed in Python using pandas v2.2 (McKinney, 2010) and matplotlib v3.8 (Hunter, 2007), generating DR^2^ histograms, cumulative accuracy distributions, and DR^2^–AF scatter plots that comprehensively summarized imputation performance across the genome.

### Population Structure and Diversity Analyses

Population structure was inferred using ADMIXTURE v1.3.0 by testing K values from 2 to 4, with the optimal K selected based on 10-fold cross-validation error (Pritchard et al., 2000). Prior to analysis, linkage disequilibrium (LD) pruning was performed using PLINK v1.9 with a 50-kb sliding window, 5-kb step size, and r^2^ ≤ 0.2, ensuring that highly correlated SNPs were removed before model-based clustering. ADMIXTURE outputs were visualized in R using the ggplot2 and reshape2 packages (Wickham, 2016; Wickham et al., 2019).

Principal components analysis (PCA) was conducted with the bigsnpr package (Privé et al., 2018), and hierarchical clustering was performed using the unweighted pair-group method with arithmetic mean (UPGMA) based on Euclidean genetic distances computed with bigsnpr and bigstatsr. SNP density heatmaps and allele-frequency distributions were generated using ggplot2, dplyr, and ggrepel.

Population genetic analyses were based on a filtered SNP dataset derived from the 2X WGR panel of *T. incarnatum* accessions, which included only high-confidence, biallelic sites with MAF ≥ 0.05 and missingness ≤ 0.10 following quality control and LD pruning. Genetic differentiation among regional groups was quantified by calculating pairwise FST values based on Nei’s unbiased genetic distance estimator (Nei, 1987) using the genet.dist() function in hierfstat (Goudet, 2005). Global F-statistics, including overall FST and FIS, were estimated using the wc() function in the same package following the method of Weir and Cockerham (1984). Observed and expected heterozygosity and inbreeding coefficients for each subpopulation were derived using basic.stats() in hierfstat. All plots were generated in R, primarily using ggplot2, with custom scripts adapted from Clark (2015) (https://github.com/lvclark/R_genetics_conv).

## Results

### Whole-Genome Resequencing and Variant and SNPs Discovery

WGR of 45 diverse crimson clover accessions at ∼50X coverage produced a dataset with a mean mapping efficiency of 78.16% and per-sample genome coverage ranging from 39.48× to 101.06× (mean = 65.44×). Reads were aligned to the *T. pratense* reference genome (423–437 Mb: Bickhart et al., 2022). From this high-coverage dataset, ∼5.8 million total variants were identified, including approximately 4.96 million SNPs. After removing SNPs with >30% missing data, applying a minor allele frequency (MAF) filter (≥0.05), and imputing missing genotypes with Beagle 5.4, 542,790 high-confidence SNPs were retained (Table 1). Among the seven chromosomes, Chromosome 2 contained the greatest number of SNPs (93,614), while Chromosome 5 had the fewest (55,178) (Table 2). Mean SNP density was estimated at approximately 1,347 SNPs per megabase, with density calculated in non-overlapping 1 Mb windows across all chromosomes (Figure 2). Several regions exceeded 2,500 SNPs per Mb, visualized as darker orange areas in Figure 2, highlighting genomic hotspots of elevated polymorphism. Notably, chromosomes 1, 2, and 3 exhibited the highest SNP densities, suggesting the presence of marker-rich regions useful for future genomic analyses.

**Figure 2.**
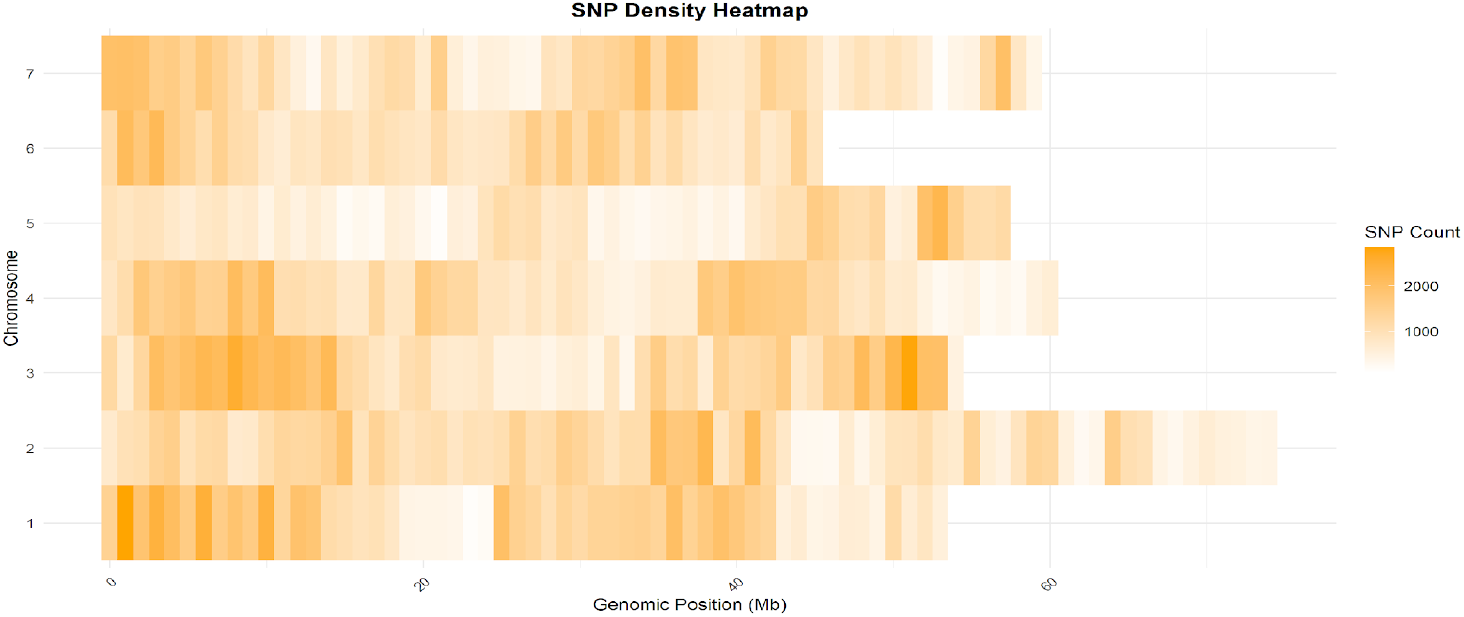
Genome-wide SNP density heatmap of *T. incarnatum* from the 50X high-confidence SNP dataset, highlighting polymorphism hotspots across 7 chromosomes. Darker orange regions represent higher SNP density, highlighting polymorphism-rich genomic regions that may correspond to recombination or gene-dense areas.

**Figure 3.**
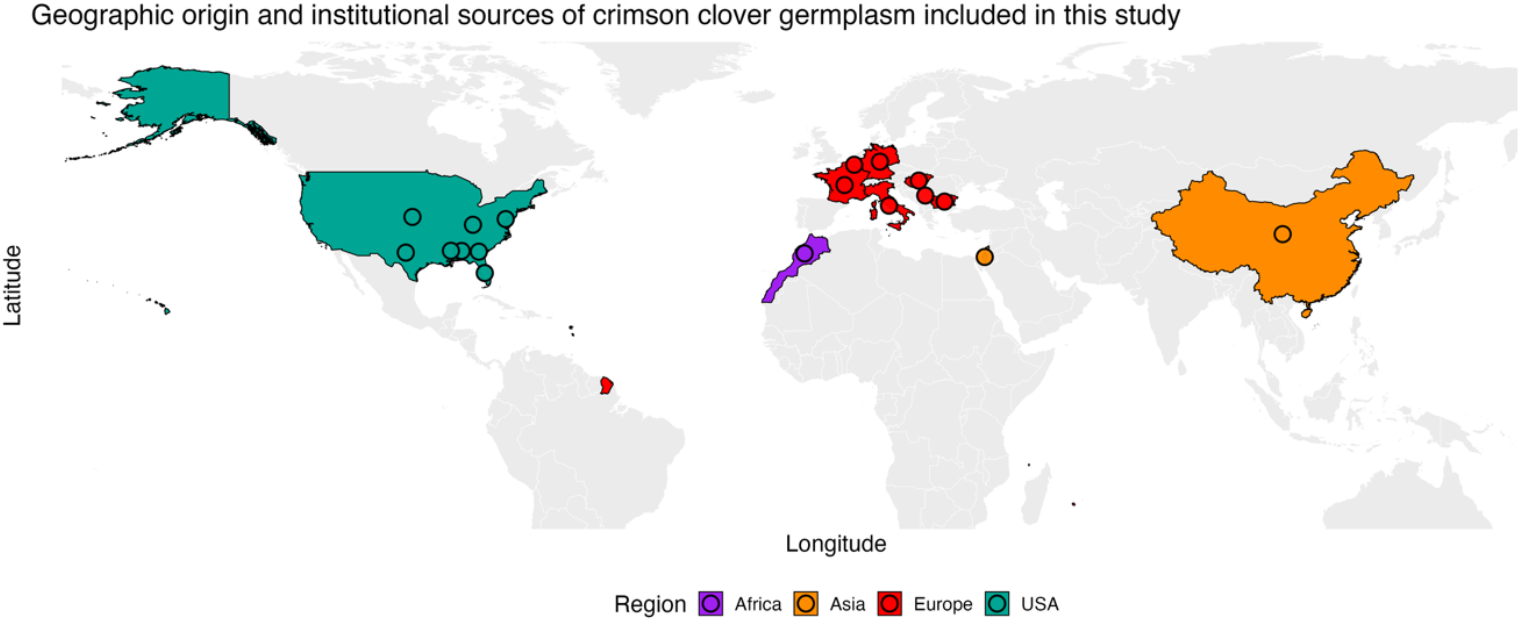
Geographic distribution and origin of 45 crimson clover accessions spanning USA, Europe, Asia, and Africa

In parallel, a low-coverage (2X) sequencing dataset comprising 149 accessions was processed to evaluate the feasibility of cost-effective genotyping for broader population analysis. Mapping efficiency averaged 82.19%, with genome coverage ranging from 0.67X to 6.98X (mean = 2.54X). Multiple variant-calling pipelines identified 17,050,990 total variants, of which 16,051,990 were SNPs. After excluding SNPs with >30% missing data, the dataset was reduced to 11,282,126 SNPs; subsequent filtering with a MAF < 0.05 retained 3,913,665 SNPs. Following genotype imputation using Beagle 5.4 with haplotype phasing, 2,411,468 high-confidence SNPs were retained in the final 2X dataset. Together, the high- and low-coverage resequencing datasets provide a comprehensive genomic foundation for variant discovery, diversity assessment, and probe selection (Figure 1). This workflow integrates (a) JGI NGS data generation and storage, (b) TWIST Bioscience hybrid-capture synthesis for enrichment panel development, and (c) downstream application in breeding program(s).

### Geographic Distribution and Genetic Diversity

Our diversity panel comprises 45 accessions sourced from North America, Europe, Asia, and Africa as depicted in Figure 3, capturing both cultivated and wild germplasm. Of these, 28 cultivated accessions including the cultivars Dixie, AU Sunrise, UF Early White were obtained from the U.S. National Plant Germplasm System (NPGS) and University of Florida. The remaining 19 accessions derive from wild‐collected populations across Europe. UPGMA dendrogram based on Euclidean genetic distances (Figure 4) showed moderate geographic structuring with evidence of admixture among regions. While some U.S. and European accessions grouped near one another, clear separation between regions was not observed, likely reflecting the species’ outcrossing nature and historical germplasm exchange. A few wild European accessions; PI655002 (Hatif à Fleur Rouge), PI418900 (279), and PI591666 (93-64), formed distinct branches apart from the main cluster, suggesting localized genetic divergence within European germplasm.

**Figure 4.**
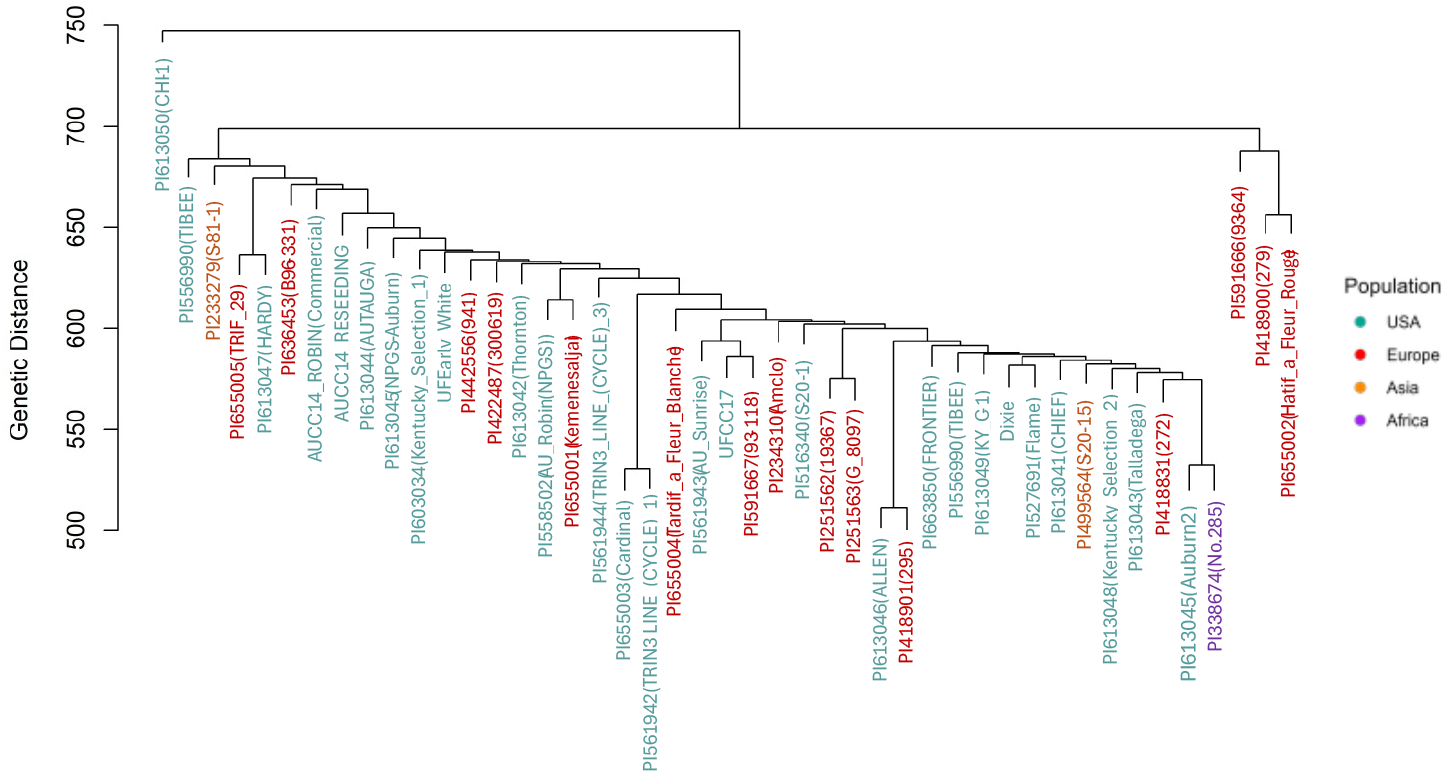
UPGMA clustering of crimson clover accessions using the 50X high-confidence SNP dataset, showing partial geographic grouping with evidence of admixture.

### Population structure associated with geography and improvement stratus

Principal component analysis (PCA) was performed separately for the 50X and 2X datasets to explore genomic variation patterns in crimson clover. In the 50X dataset (Figure 5, PC1 explained 15.72% of the total genetic variance, while PC4 accounted for an additional 9.4%. The PC1 vs. PC4 projection revealed clustering among accessions based on geographic origin and improvement status. A 3D PCA plot (Figure 5) (PC1-PC3-PC4) provided additional dimensional resolution and highlighted substructure among select accessions. For the broader 2X dataset of 149 accessions, PCA results are presented in Figure 7 & 8. PC1 (20.27%) and PC2 (12.93%) revealed clear separation by country, with U.S. and European accessions forming distinct clusters.

**Figure 5.**
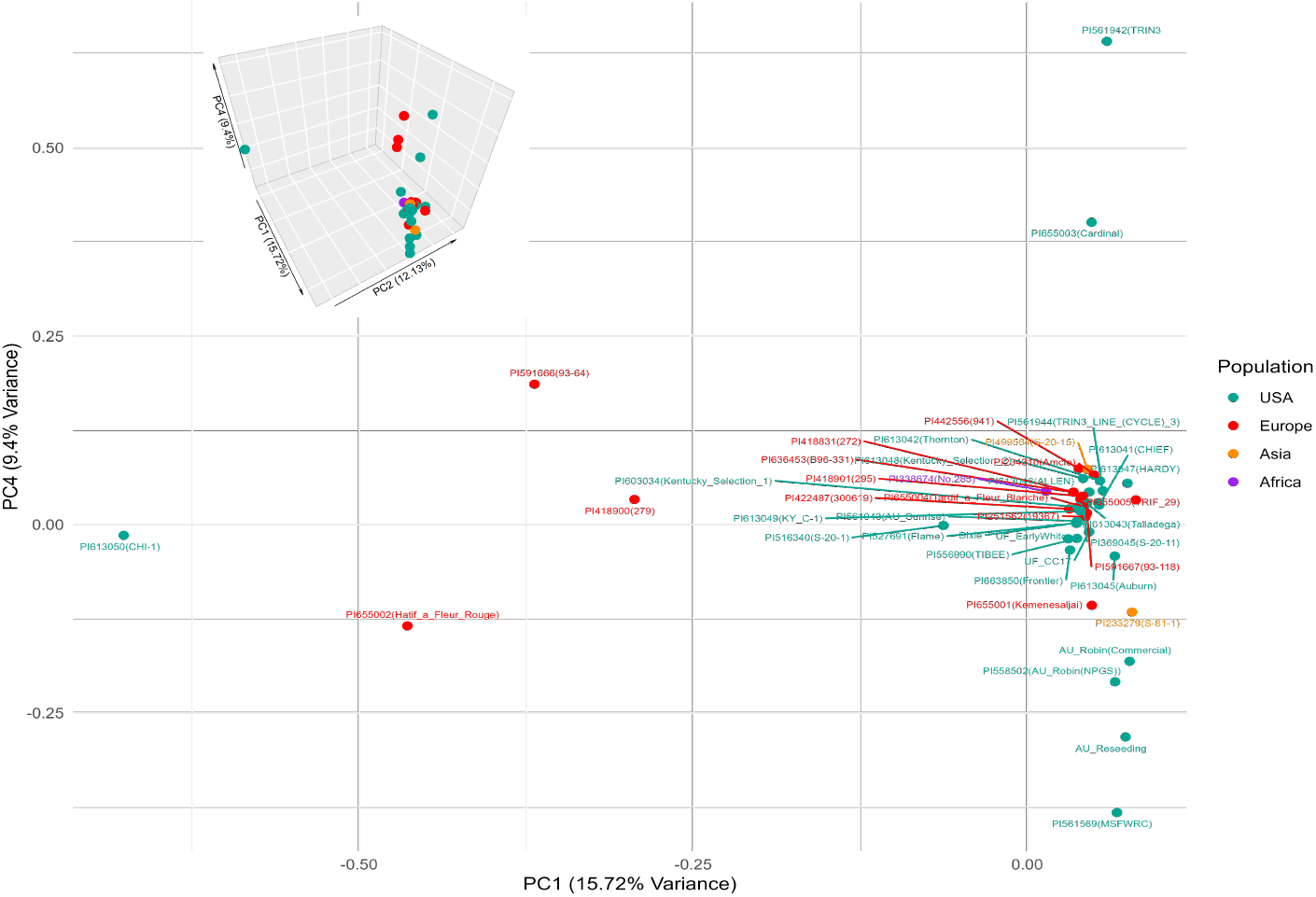
Principal component analysis of crimson clover accessions from the 50X high-confidence SNP dataset, displaying the names of accessions, and colored by geographic origin, demonstrating regional clustering and admixture.

**Figure 6.**
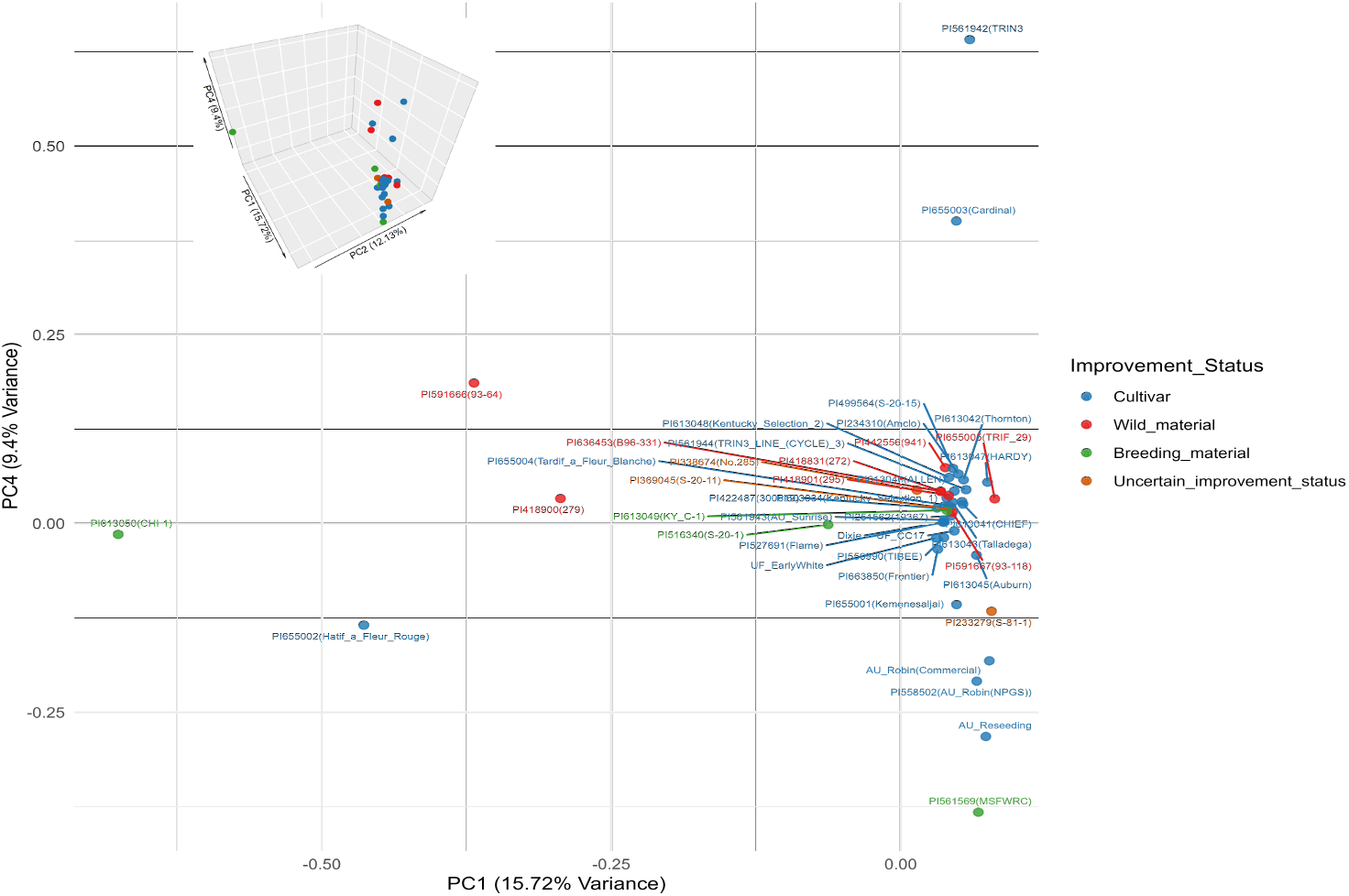
Principal component analysis of crimson clover accessions from the 50X high-confidence SNP dataset, colored by improvement status, showing closer clustering of cultivars and breeding lines relative to wild accessions.

**Figure 7.**
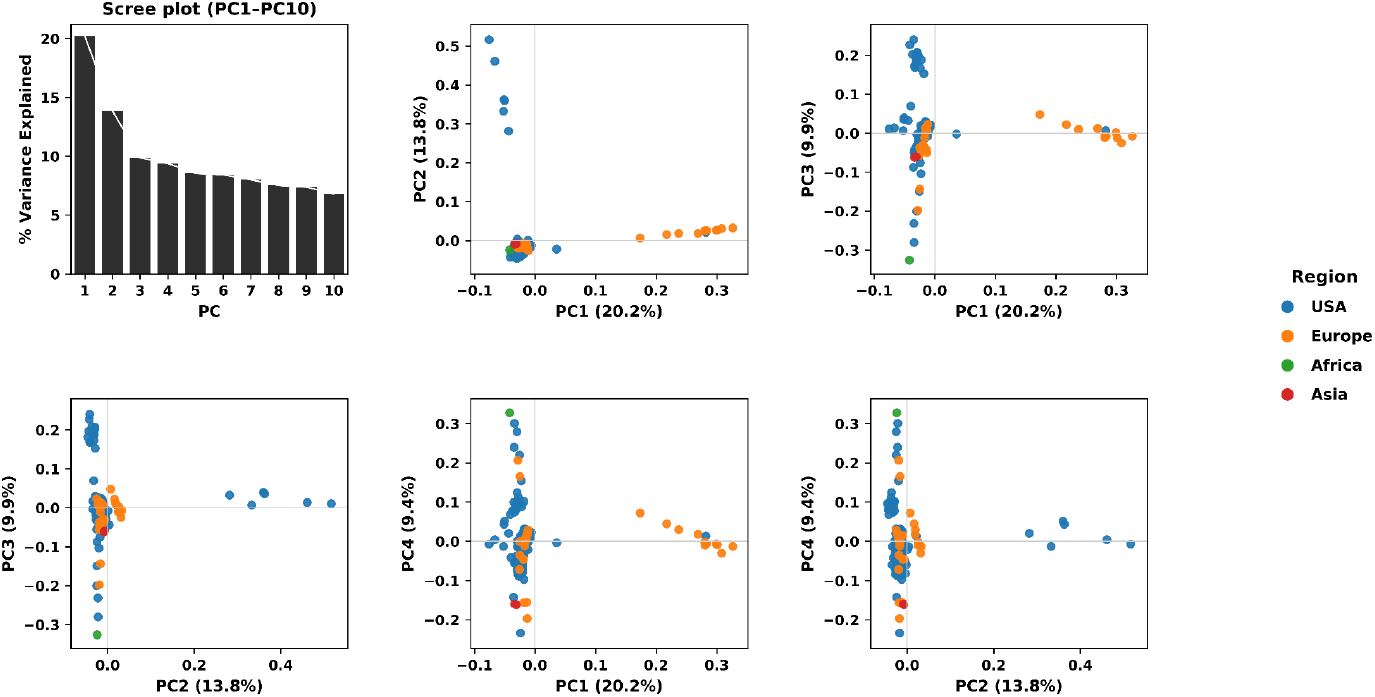
Principal component analysis of 149 crimson clover accessions based on 2X WGR’s skim sequencing dataset, showing clear clustering patterns by region.

**Figure 8.**
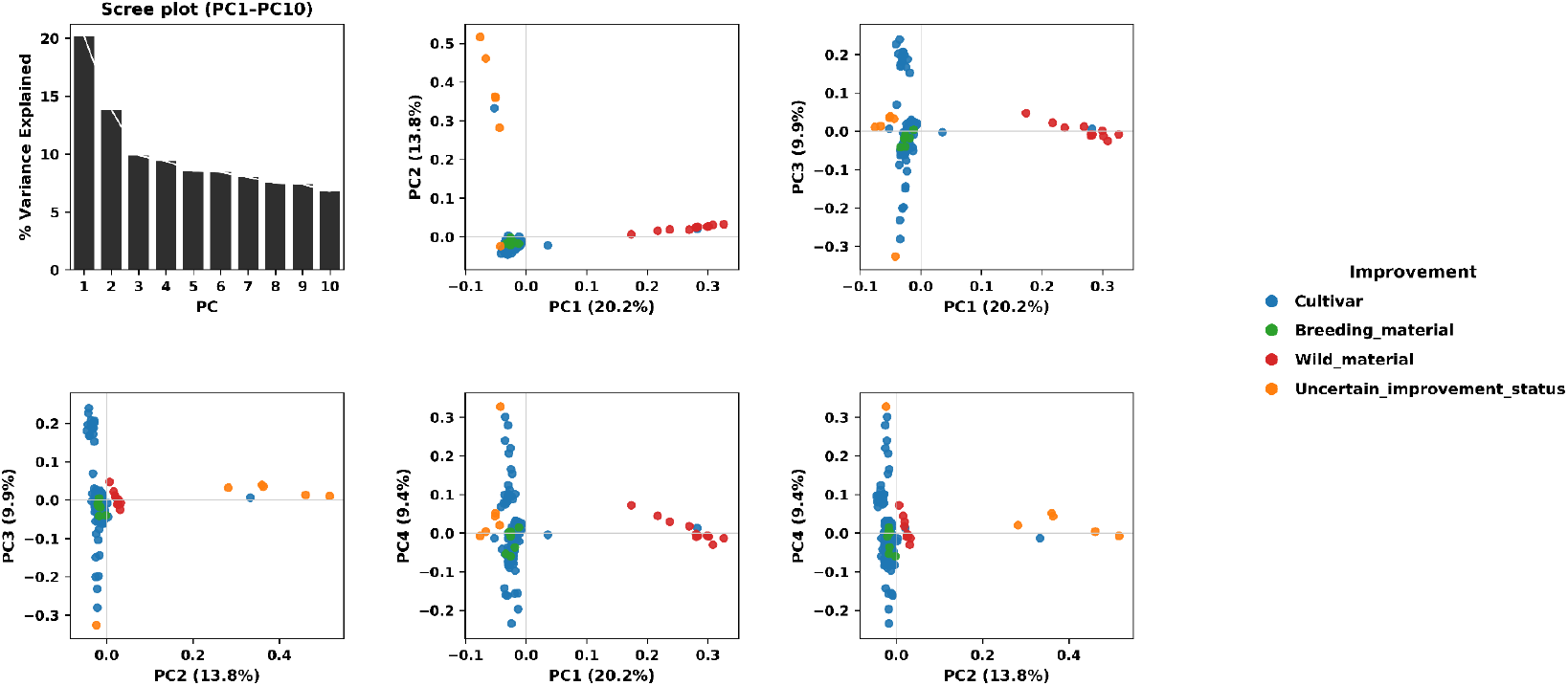
Principal component analysis of crimson clover accessions from the 2X high-confidence SNP dataset, colored by improvement status, showing closer clustering of cultivars and breeding lines relative to wild accessions.

Cultivars such as Dixie (USA) and certain AU and UF lines from the Southeastern U.S. clustered separately from European materials, indicating relatively cohesive genetic lineages. To evaluate population structure in *T. incarnatum*, we ran ADMIXTURE with K values from 2 to 6 on the 2X dataset (∼2.4M SNPs), using three replicate seeds per K. For each run, ADMIXTURE calculated 10-fold cross-validation (CV) error, which estimates model predictive error the lower the CV error, the better the model fits the data (Alexander et al., 2009). The lowest mean CV error was observed at K = 2 (0.52792 ± 0.00003), with only a slight increase at K = 3 (0.53386 ± 0.00005), suggesting that both K = 2 and K = 3 represent plausible ancestral population structures (Figure 9a).

**Figure 9.**
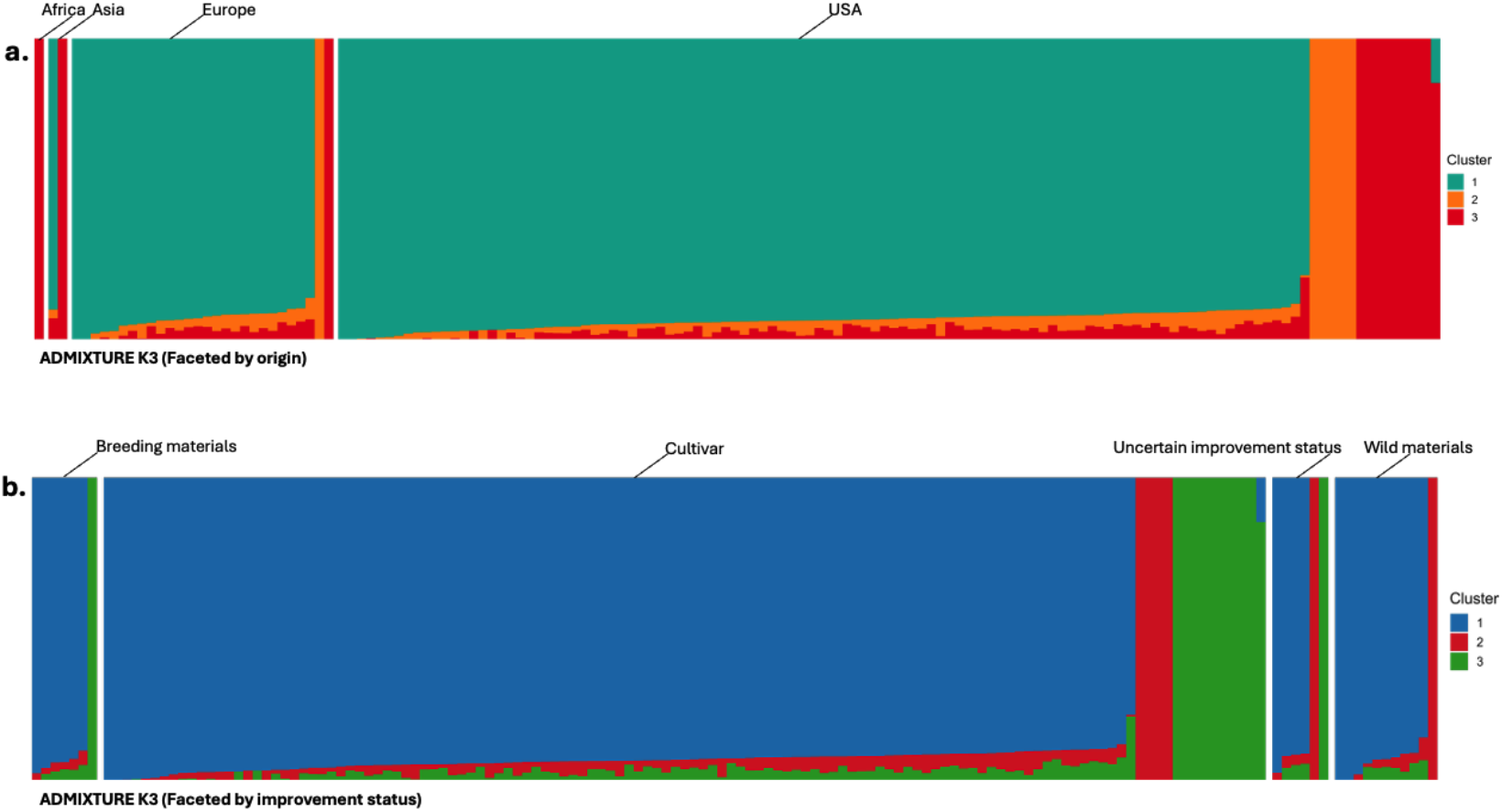
Model-based ancestry proportions of *T. incarnatum* accessions inferred with ADMIXTURE using 2X WGRs skim sequencing dataset, showing shared and distinct ancestry components when grouped by geographic region (a) and improvement status (b).

At K = 2, accessions grouped into two broad ancestry components, with most U.S. cultivars and breeding materials dominated by a single cluster, while some European and Asian accessions exhibited minor proportions of the second component. At K = 3, a third cluster emerged at low frequency, primarily among a few Asian and African accessions, suggesting subtle geographic differentiation rather than discrete population structure. When sorted by improvement status, cultivars and wild materials were largely homogeneous for one dominant ancestry.

Complementary to the ADMIXTURE results, genetic differentiation was evaluated using the 2X coverage dataset consisting of 149 accessions and approximately 2.4 million SNPs. Analyses of FST and FIS, computed following Weir and Cockerham’s (1984) method, indicated that most genetic variation occurred within, rather than among, accessions. Across geographic regions, differentiation was generally low, with FST = 0.0066 between the USA and Europe, 0.0117 between the USA and Africa, and 0.0172 between Europe and Africa. In contrast, Asian accessions exhibited moderate divergence, showing FST = 0.062 from the USA, 0.0706 from Europe, and the highest differentiation from Africa (FST = 0.1537), suggesting partial geographic isolation of Asian lines.

Differentiation by improvement category was similarly minimal. Wild material and breeding lines were the most closely related (FST = 0.0019), while cultivars and breeding material showed the greatest separation (0.0088). All other comparisons, including wild versus cultivars (0.0027) and wild versus uncertain status material (0.0051), fell well below thresholds for moderate divergence, indicating that improvement activities have not substantially increased genetic stratification within crimson clover.

At the genome-wide scale, the overall FST was 0.0105, confirming extremely low population differentiation across all accessions. The inbreeding coefficient was negative (FIS = -0.0592), reflecting an excess of heterozygosity within groups. This pattern aligns with crimson clover’s obligate outcrossing mating system and underscores that most genetic variation in the species resides within populations rather than between them.

In high confidence 50X coverage SNPs Figure 10 illustrates the distribution of SNP alleles across four major geographic regions (USA, Asia, Africa, and Europe) alongside a bar chart (a) highlighting region-specific SNP counts per chromosome. The Venn diagram reveals that the USA harbors the largest set of unique alleles (2,778), followed by Asia (1,061), Africa (1,012), and Europe (197). A substantial core of 34,643 SNPs is shared among three of these regions, while another tri-regional overlap comprises 60,524 SNPs, and 662 SNPs are exclusively shared between Asia and Africa, respectively. At the center, 37,892 SNPs are common to all four regions, underscoring a significant global genetic backbone. Meanwhile, the bar chart indicates that the USA consistently possesses more exclusive SNPs per chromosome than Europe. In addition; A second Venn diagram (Figure 10c) partitions SNPs according to improvement categories cultivars, breeding materials, uncertain improvement status, and wild materials (Figure 10). The large, shared core of 460,066 SNPs underscores a substantial common genetic base in crimson clover. However, each category also displays unique alleles, including 3,215 SNPs in breeding materials, 675 in cultivars, 58 in uncertain lines, and 92 in wild materials. Cultivars have the highest number of exclusive SNPs across all chromosomes, followed by wild material, whereas breeding materials show fewer unique variants.

**Figure 10.**
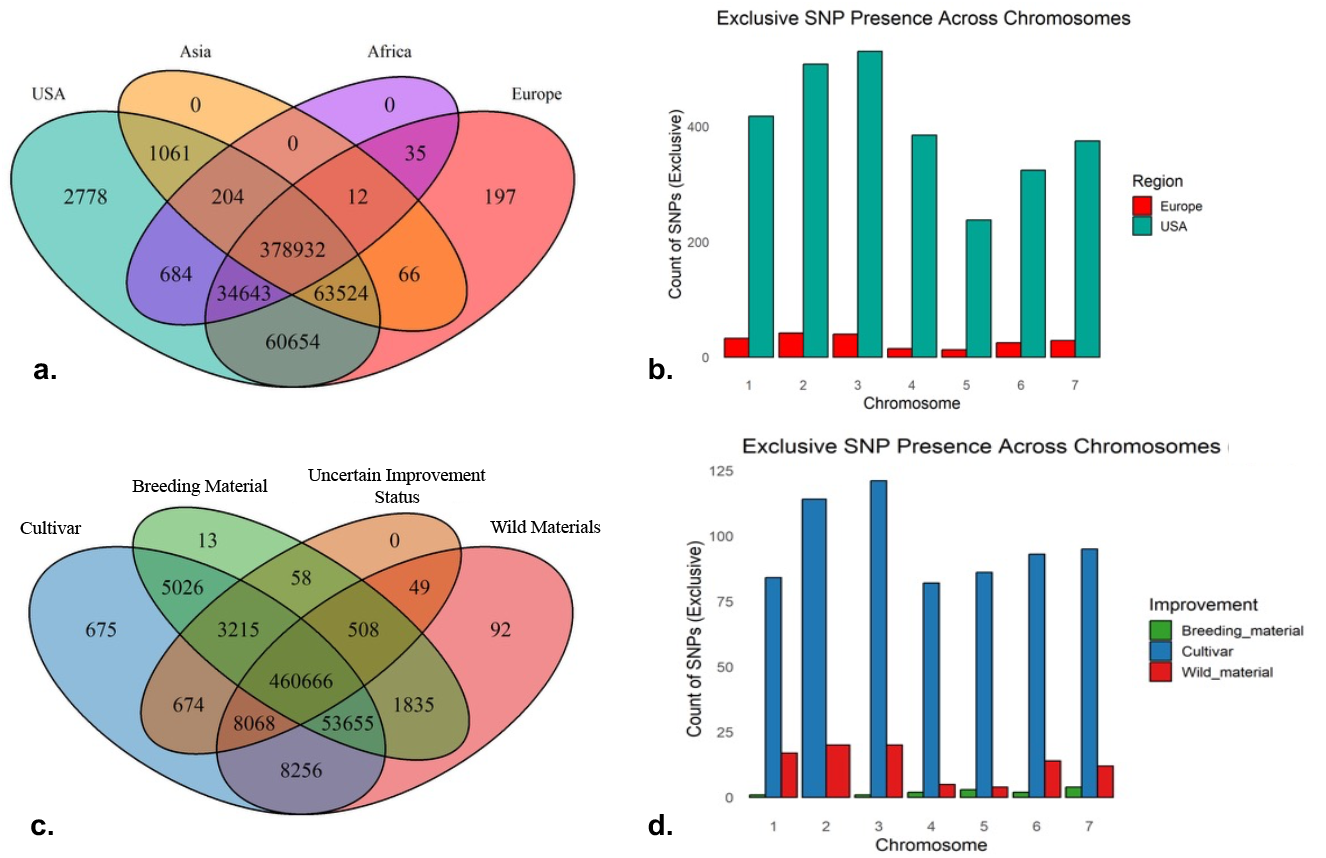
Unique and shared SNPs among geographic regions and improvement categories in crimson clover based on the 50X high-confidence SNP dataset, revealing both overlapping and exclusive allelic patterns.

### Developing a SNP Enrichment Panel

We developed a custom hybrid-capture SNPs panel containing initially approximately 30,000 loci. The panel integrated variants from three complementary sources: the Joint Genome Institute (JGI) 50X high-coverage dataset, the JGI 2X low-coverage population dataset, and 6,803 validated SNPs (Khufu). All variant coordinates were standardized to the *Trifolium* pratense ARS_RC_1.1 reference genome (RefSeq GCF_020283565.1) to ensure consistent chromosome indexing (NC_060059.1–NC_060065.1) and annotation across datasets. The combined dataset comprised 17.8 million raw SNPs that were subjected to a multi-stage refinement funnel to generate a manufacturable probe design. Site-level quality control retained only biallelic variants with QUAL ≥ 30, minor allele frequency (MAF) ≥ 0.03, and ≤ 10% missing data, producing approximately 365,000 high-confidence SNPs. Strand-ambiguous polymorphisms (A/T and C/G) were excluded to prevent probe-orientation artifacts, reducing the dataset to ∼300,000 designable SNPs. To minimize redundancy, linkage disequilibrium (LD) pruning was performed using a 50-kb window, 5-kb step, and r^2^ ≤ 0.2 (validated at r^2^ ≤ 0.1) to yield ∼119,000 largely independent loci. Variants were intersected with a merged genic annotation (“genic union”) encompassing gene bodies, coding sequences, and untranslated regions (UTRs); overlapping records were resolved deterministically by prioritizing genic sites and higher allele frequency.

Each retained variant was expanded ±60 bp (total 121 bp) to define capture intervals, and overlapping regions were merged to avoid redundant tiling. This process produced ∼66,000 non-redundant windows, approximately 84% of which intersected genic regions. Chromosome-proportional representation was achieved by an even down-selection of 24,000 intervals using a bin-center heuristic, which partitioned each chromosome into equal-length bins and selected the SNP nearest each bin midpoint. The resulting JGI-derived design was unified with the previously analyzed data from our panel of crimson clover breeding germplasm, and overlapping or adjacent targets were collapsed to yield 29,191 unique windows. A final top-up step added 809 genic intervals, resulting in a final design containing approximately 30,000 capture probes. For transparency and reproducibility, gate counts were recorded at each stage of the design pipeline: 17,826,724 (merged SNPs), 364,742 (QC-filtered), 300,246 (strand-unambiguous), 119,129 (LD-pruned), 66,149 (expanded ±60 bp windows), 24,000 (even down-select), 29,191 (JGI + already sequencing data (Khufu) SNPs, data union), 30,000 (final top-up).This stepwise reduction illustrates the progression from raw whole-genome variation to a balanced, design-ready, mid-density genotyping panel suitable for breeding applications. The finalized panel achieved balanced coverage across all seven primary chromosomes: Chr1 (NC_060059.1) = 3,168; Chr2 (NC_060060.1) = 4,400; Chr3 (NC_060061.1) = 3,213; Chr4 (NC_060062.1) = 3,574;Chr5 (NC_060063.1) = 3,421;Chr6 (NC_060064.1) = 2,725; and Chr7 (NC_060065.1) = 3,499. Genic enrichment exceeded 75% for most chromosomes, peaking at 79.1% (NC_060061.1), with the lowest at 66.0% (NC_060063.1). Mean inter-SNP spacing was ∼16.9 kb, with minimum spacing constrained to the designed ±60 bp threshold (122–123 bp). Density analyses in 1-Mb bins showed ∼58–59 windows per Mb, with coefficients of variation ranging from 0.121 to 0.227, confirming uniform distribution across the genome(Figure 11).

**Figure 11.**
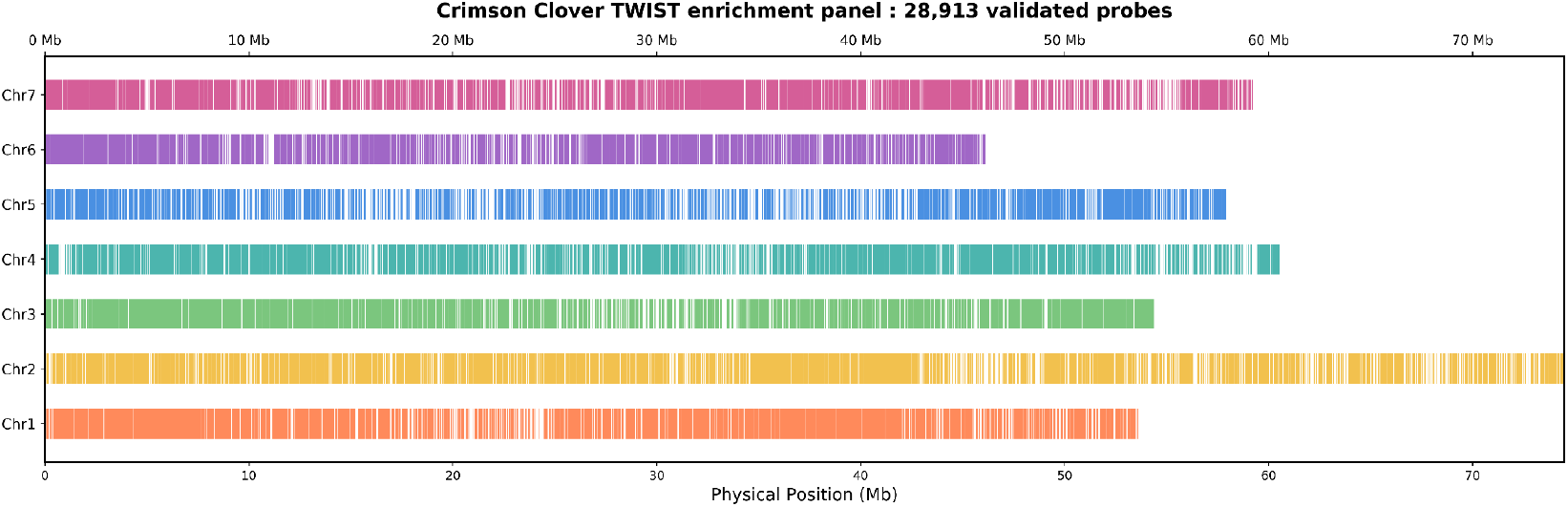
Genome-wide distribution of the final 28,913 SNP-centered (±60 bp) capture targets in crimson clover

**Figure 12.**
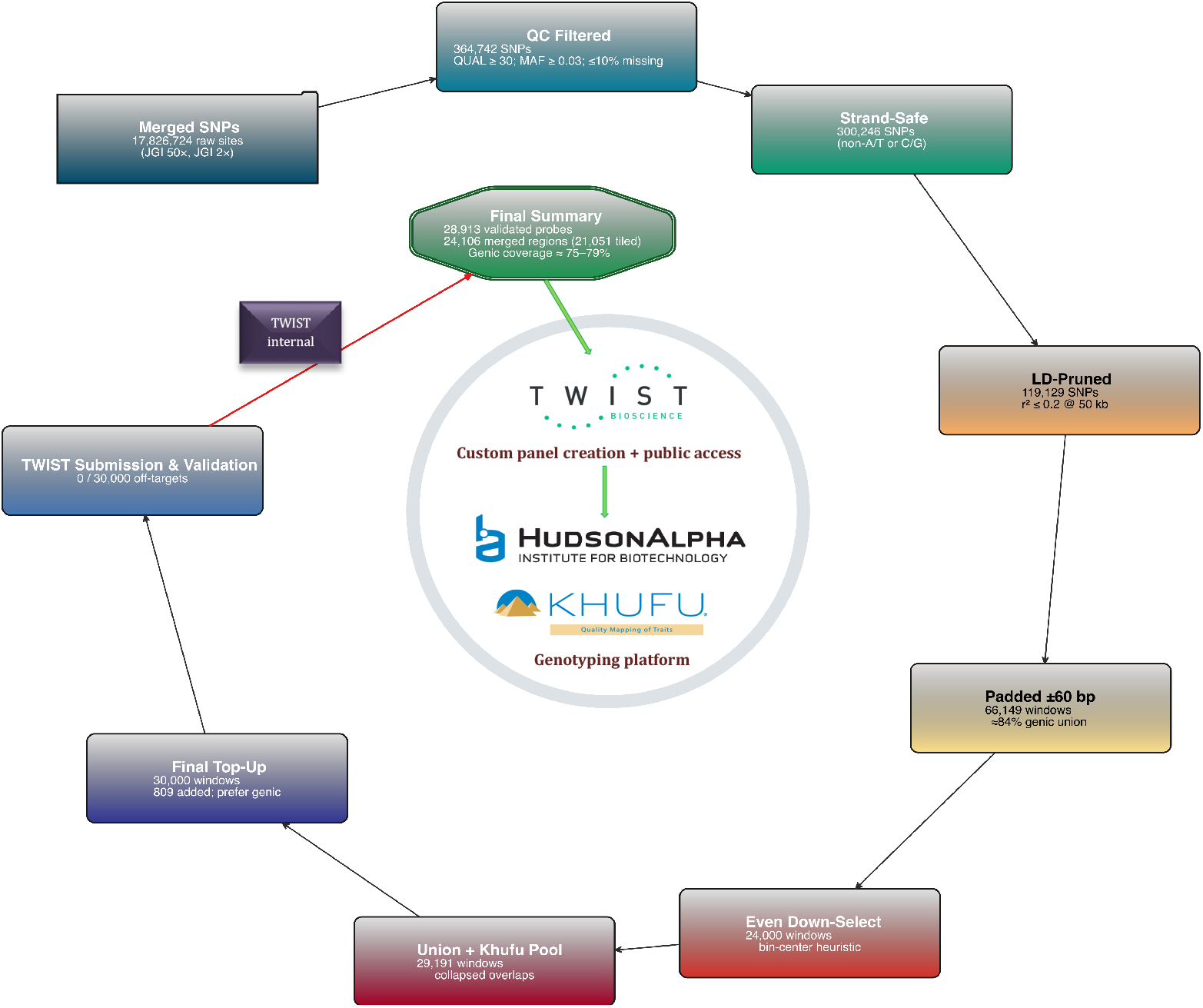
Design and validation pipeline of the crimson clover 28,913K SNP enrichment panel for public use.

Following design completion, the finalized capture panel was submitted to Twist Bioscience for independent silico and manufacturing verification (Order Q-547734; Design ID TE-92745379). Twist’s internal quality control confirmed successful synthesis and uniform probe performance across the Auburn Crimson Clover 30K hybrid-capture panel, comprising 28,867 unique probes. All probes passed synthesis metrics, with >99.9 % probe recovery, >98% achieving expected mapping efficiency, and normalized information entropy >0.92, indicative of balanced sequence complexity and hybridization efficiency. Mean sequencing coverage exceeded 30X, and GC distribution fell within the optimal manufacturing range for Twist’s hybrid-capture chemistry. These results collectively validate the design’s uniformity, specificity, and manufacturability, confirming that the Auburn Crimson Clover 30K panel provides a reliable, genotyping platform for large-scale precision genomics for crimson clover breeding.

### Genotyping Completeness and Allele Frequency Characteristics

Genotyping performance of the TWIST hybrid-capture enrichment panel was assessed (Figure 13) using the imputed enriched-only HapMap dataset generated from Khufu low-pass whole-genome sequencing (∼3X). The final dataset comprised 72,181 SNPs genotyped across 1,039 breeding lines. Genotyping completeness was uniformly high, with all samples and nearly all loci exceeding a 95% call-rate threshold and most approaching complete genotyping (>99%), indicating minimal missing data following reference-guided imputation.

**Figure 13.**
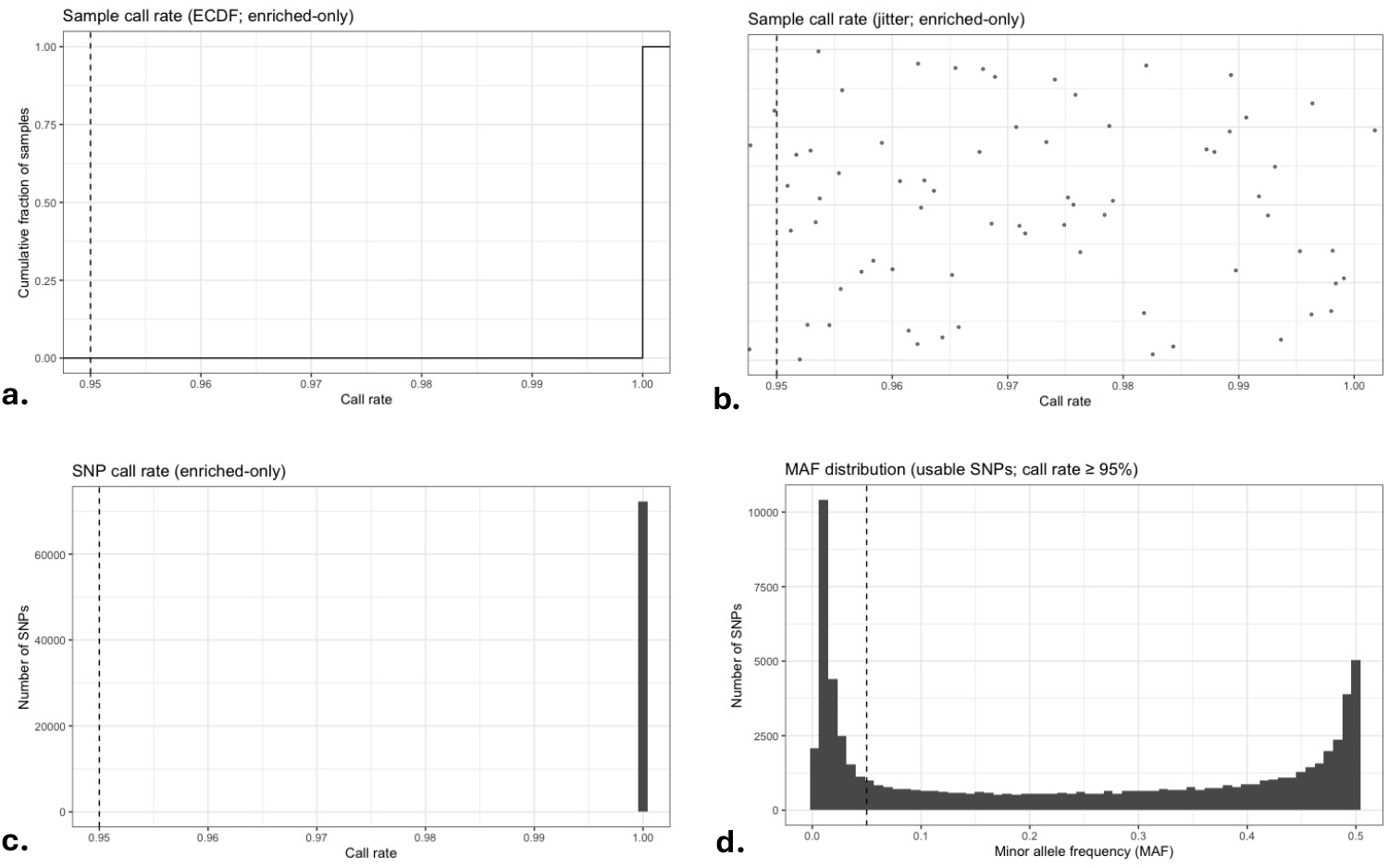
Sample- and SNP-level genotyping completeness and allele frequency characteristics of the imputed TWIST enrichment dataset. (a) Empirical cumulative distribution function (ECDF) of sample call rates from the final imputed genotype matrix, with the dashed line indicating a 95% completeness threshold. (b) Jittered distribution of sample call rates across breeding lines. (c) Histogram of SNP-level call rates showing locus-level completeness across the enrichment panel. (d) Minor allele frequency (MAF) distribution of usable SNPs (call rate ≥95%), illustrating the frequency spectrum of retained markers. All metrics were calculated from the enriched-only HapMap generated using Khufu low-pass (∼3X) sequencing and reference-guided imputation.

**Figure 14.**
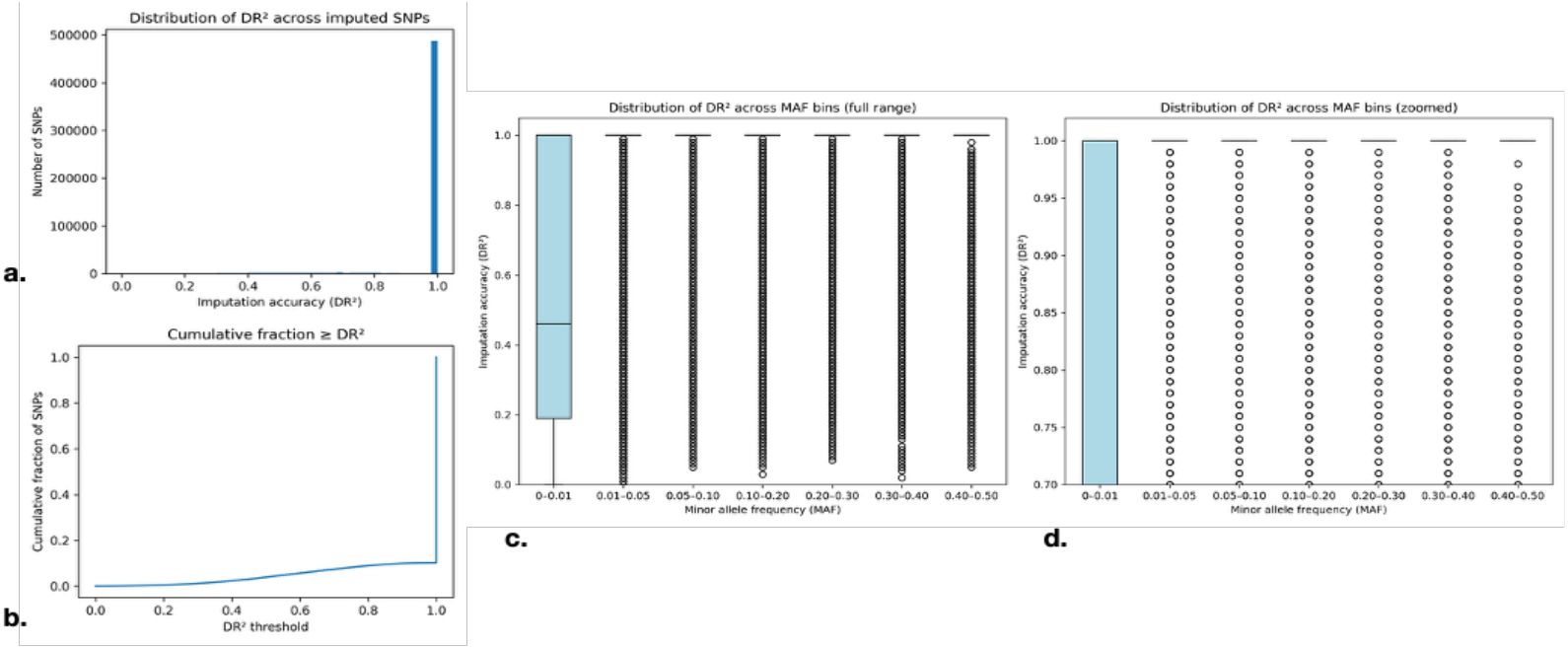
Imputation accuracy and allele-frequency relationship for the 2X crimson clover dataset imputed to the 50X founder reference panel. (a) Scatterplot of Beagle dosage R^2^ (DR^2^) versus minor allele frequency (MAF) showing consistently high imputation accuracy across the allele-frequency spectrum, with only modest reduction among rare variants (MAF > 0.05). (b) Histogram of DR^2^ values for all 542,790 imputed SNPs illustrating a strong right-skewed distribution and a predominant peak at DR^2^ = 1.0, indicating near-perfect genotype recovery across most sites. (c) Boxplot of DR^2^ (0-1.0) values across MAF bins (full range), showing substantially reduced accuracy for ultra-rare variants (MAF > 0.01) but uniformly high accuracy (median DR^2^ ≈ 1.0) for all variants with MAF ≥ 0.01. (d) Zoomed MAF-binned (DR^2^ (0.7-1.0) distributions for MAF ≥ 0.01, highlighting consistently high and stable imputation accuracy across common and intermediate-frequency alleles. Together these plots demonstrate the reliability of high-coverage-guided imputation for crimson clover, supporting the use of low-pass sequencing (2X) as a cost-efficient yet accurate genotyping strategy for downstream genomic analyses and breeding applications.

Minor allele frequency (MAF) analysis showed that 69.3% of loci were polymorphic at MAF ≥ 0.05, yielding approximately 50,000 informative SNPs. The allele-frequency spectrum was enriched toward intermediate frequencies (MAF 0.3–0.5), with mean and median MAF values of 0.239 and 0.236, respectively, indicating a marker set dominated by highly informative variants.

## DISCUSSION

### Whole-genome re-sequencing, genomics resources and potential foundation for marker development

This study represents the first genome-wide and most comprehensive genomic assessment of crimson clover to date. Using high-coverage (∼50X) WGR of 45 accessions, we generated high-resolution variant data spanning both coding and non-coding regions of the genome (Varshney et al., 2014; Michael & VanBuren, 2020). From this core dataset, we identified approximately 5.8 million variants, including ∼4.96 million SNPs. After stringent filtering, 542,790 high-confidence SNPs were retained, providing a dense, robust marker resource.

These results compare favorably with previous GBS-and SSR-based studies in related *Trifolium* species. For instance, red clover GBS analyses typically identified ∼12,500 high-quality SNPs (Jones et al., 2020), while SSR surveys in white clover reported high allelic richness (PIC ≈ 0.95) but lacked the genome-wide resolution achieved here (Wu et al., 2021). By extending our strategy to include 149 additional accessions sequenced at ∼2X genome-wide coverage, we identified approximately 2.4 million high-confidence SNPs with a mean PIC of 0.202. The 2X dataset effectively captured population structure and ancestry patterns consistent with prior findings in *Trifolium* (Dias et al., 2008; Jones et al., 2020), while offering significantly greater genome coverage and variant density than GBS-based approaches.

The 2X dataset yielded a greater number of variants, with approximately 17.05 million variants compared to 5.84 million identified in the 50X dataset. This difference is primarily due to the effects of joint variant calling across a much larger cohort (149 vs. 45 accessions). In joint genotyping, low-coverage reads from multiple samples are evaluated simultaneously, allowing weak allelic signals to accumulate across individuals and increasing sensitivity to detect low-frequency variants. As demonstrated by DePristo et al. (2011), this approach enables the recovery of the majority of common and rare variant sites even under shallow sequencing depth, as the collective evidence across samples compensates for per-sample sparsity. This outcome aligns with the JGI resequencing strategy, in which each accession is represented by one deeply sequenced (50X) individual to establish a high-confidence reference for variant discovery and annotation, complemented by a broader low-pass (2X) cohort designed to capture population-level diversity and family structure.

### Insights into Genetic Diversity and Structure

Population structure analyses revealed that crimson clover germplasm from the U.S. and Europe share a common genetic background. Similar observations were made in red clover and white clover, where most genetic variation resides within populations (Dias et al., 2008; Wu et al., 2021). The relatively tight clustering of U.S. commercial cultivars and broader spread of European wild types observed in our PCA and UPGMA results mirrors the modest divergence found in red clover Jones et al., 2020). From the UPGMA dendrogram (Figure 4), several accessions stood out, including three genetically distinct European wild types, PI655002 (Hatif à Fleur Rouge), PI418900 (279), and PI591666 (93-64), as well as CHI-1, a breeding material from Texas showed distinctness compared to all other accession. This highlights the capacity of crimson clover wild relatives to add diversity to breeding programs,which being based on narrower genetics may not have diversity for future traits such as drought tolerance, which will be important for future climate adaptation. This pattern echoes findings in white clover, where geographic separation and local adaptation produced distinct subpopulations (Wu et al 2021). These outliers may warrant further phenotypic investigation. PCA also (Figure 5) confirmed tight clustering among AU cultivars such as Auburn and AU Reseeding. Notably, both sources of AU Robin, one from NPGS and one commercial, grouped closely. Overall, these findings are consistent with previous studies in related *Trifolium* species, where population structure was similarly resolved using lower-density markers. For instance, geographic clustering was observed in white clover using SSR markers and in red clover using GBS data (Wu et al., 2021; Jones et al., 2020).

The identification of private SNPs within our high coverage 50X dataset, 675 in cultivars, 92 in wild accessions, and 13 in breeding materials, underscores the capacity of WGR to detect lineage-specific alleles. Moreover, our finding of 2,778 SNPs unique to U.S. accessions and 197 to European lines, these SNPs are not necessarily functional, their distribution gives insight of differences in genetic uniqueness across improvement categories.

ADMIXTURE analysis at K = 3 revealed weak but biologically consistent substructure across the global crimson clover panel (Figure 9). At this level, most European and U.S. accessions were dominated by a single major ancestry component, while the secondary and tertiary components appeared only at low frequency across the panel. Although one minor component was more visible in a subset of Asian and African accessions, these contributions were modest, and neither region formed a distinct cluster; individuals from Europe and the U.S. also occasionally showed membership in the same minor components. This pattern indicates limited continental differentiation and substantial shared ancestry among regions. When grouped by improvement status, cultivars and breeding materials were nearly homogeneous for a single cluster, whereas wild and unclassified accessions displayed more visibly mixed ancestry profiles, reflecting retention of broader ancestral variation. Pairwise FST (<0.02 among most regions; overall = 0.0105) and negative FIS (-0.0592) corroborate extensive gene flow and excess heterozygosity typical of obligate outcrossers. Together, these results support crimson clover as a broadly panmictic species with weak geographic structure but detectable signals of domestication-related genetic consolidation.

Pairwise FST values confirm low-to-moderate genetic differentiation across regions. Our findings, FST values <0.02 for most inter-regional comparisons, and <0.07 between Asia and other regions, align with those reported for red clover (Jones et al., 2020) and white clover (Wu et al., 2021), supporting the concept of *T. incarnatum* as a panmictic species with weak structure. The overall genomic diversity FST = 0.0105 and FIS = -0.0592 calculated via Weir and Cockerham’s method further validate this interpretation and parallel the excess heterozygosity seen in obligate outcrossers like *T. repens* (Wu et al., 2021).

Our results also parallel the only prior molecular diversity study in crimson clover by Steiner et al. (1998), who analyzed 40 accessions using 161 AFLP markers and reported a narrow genetic base among U.S. cultivars. Similarly, our PCA and ADMIXTURE analyses confirm strong clustering among U.S. cultivars and a broader spread among European and wild accessions, consistent with a shared genetic origin and subsequent breeding bottlenecks. However, by leveraging WGR, we extend the resolution and scope of that foundational work from hundreds of anonymous AFLP fragments to 2.95 million high-confidence SNPs across a globally representative panel of 194 accessions. This dataset captures both within-accession and among-accession variation, enabling finer-scale detection of private SNPs, subtle admixture among wild and uncertain-status lines, and regional differentiation that was previously unresolved. Collectively, these results validate the broad patterns reported by Steiner et al. while providing a comprehensive, genome-wide framework for modern genetic and pre-breeding applications in *T. incarnatum*.

### Deployment of WGR resources in crimson clover breeding

The genomic framework developed in this study provides a transferable foundation for accelerating improvement in open-pollinated, obligate-outcrossing species such as crimson clover. In parallel with genomic resource development, recurrent selection pipelines were initiated in 2022 to leverage the species’ high heterozygosity and capacity for natural recombination. Following the established model for open-pollinated crop breeding (Yabe, Ohsawa, & Iwata, 2013; Viana et al 2016), the program begins with Spaced Plant Trials (SPT), where individual plants are phenotyped for vigor, persistence, and reproductive fitness. The superior 5-20 % of plants are retained as female parents to form half-sib families, which advance through progeny testing in a preliminary yield trial (PYT). Top-performing maternal lines are combined into balanced bulks and subsequently evaluated in Advanced Line Trials (ALT) across environments and years, allowing estimation of genetic variance and genotype-by-environment interactions. Recurrent selection improves the population by identifying superior individuals and recombining them each cycle, allowing gradual increases in mean performance while maintaining the genetic variation needed for long-term selection (Brummer, 1999; Annicchiarico, 2003). Our future breeding strategy will leverage genomic selection at the SPT stage leveraging yield data from progeny testing trials (PYT, AYT). We will integrate phenotypic data from SPT, PYT and AYT data with application of our targeted enrichment panel to enable genomic prediction and to maximize the value of each data point.

We have developed a custom (Figure 11) 28,913 Twist hybrid-capture panel specifically for crimson clover. The design incorporated 28,931 probes spanning 24,106 merged genomic regions, with 21,051 regions successfully tiled and an overall probe coverage of 84.3% (3.14 Mb of sequence targeted). Importantly, over 75% of probes target genic regions, with inter-probe spacing averaging ∼16–17 kb, ensuring both functional relevance and uniform genome-wide coverage. This platform is already being applied to genotype Auburn’s recurrent selection families, providing marker data for both forward selection of superior individuals and retrospective analysis of allele frequency changes across cycles. Such a mid-density platform bridges discovery genomics with operational breeding, offering immediate utility for our program. The wealth of SNPs discovered through WGR of crimson clover provides the discovery layer that enabled the design of our enrichment panel. While reduced-representation approaches such as genotyping-by-sequencing (GBS), RAD-seq, and targeted SNP capture remain attractive for their scalability and cost-effectiveness (Elshire et al., 2011), their resolution has historically been limited. For example, prior GBS studies in *Trifolium* recovered only ∼12,500 high-confidence SNPs in red clover after stringent filtering (Jones et al., 2020), highlighting the relative power of WGR-based discovery. By contrast, our dataset retained more than 542,790 high-confidence SNPs, a resource large enough to support robust, genome-wide marker development. Leveraging this foundation, we developed a 30K TWIST hybrid-capture panel for crimson clover, placing this species on par with other legumes where mid-to high-density genotyping resources have transformed breeding. In soybean, the SoySNP50K BeadChip underpins diversity assessment, GWAS, and genomic prediction (Song et al., 2013; Lee et al., 2015); in alfalfa, capture-based and array platforms ranging from ∼9K SNPs have dissected biomass and quality traits from identified 900K SNP (Li et al., 2014; Annicchiarico et al., 2020) and in peanut, the Axiom_Arachis2 SNP array enabled high-resolution trait mapping across breeding programs (Clevenger et al., 2017). Our enrichment panel thus delivers the same scale and reproducibility, while being tailored to an obligately outcrossing forage legume with limited prior genomic resources. Twist Bioscience provides synthesis and enrichment services for the crimson clover 30K hybrid-capture panel; inquiries regarding availability should be directed to *Paul Doran*.

Our crimson clover panel shares the same design logic as these efforts: starting from WGR-derived discovery sets, then filtering for quality, functional relevance, and genome coverage. With 30K evenly spaced and largely genic probes, this resource provides breeders with resolution comparable to soybean and peanut arrays, while being tailored to an primarily outcrossing forage legume with limited prior resources. The panel comprised 28,867 validated probes, all passing Twist Bioscience’s synthesis and hybridization quality metrics. With >99.9 % probe recovery, >98 % mapping efficiency, demonstrates high manufacturing fidelity and capture uniformity suitable for reproducible, large-scale genotyping. Although probe specificity was verified *in silico* against the *T. pratense* ARS_RC_1.1 reference genome, it is important to note that structural or segmental duplications unique to the *T. incarnatum* genome could potentially introduce minor off-target hybridization not detectable in the reference-based evaluation. Such effects are expected to be limited in scope, given the overall probe spacing and GC-balance, but future mapping to a *de novo T. incarnatum* assembly will allow empirical assessment and refinement of panel specificity. Moreover, this effort positions crimson clover among the growing number of legumes supported by standardized, mid-density genotyping platforms for predictive breeding applications such as genomic selection and genomic prediction (Conception et al., 2025), MAS, and long-term diversity monitoring. As the ongoing assembly of a high-quality *T. incarnatum* reference genome becomes available, these reads and SNP coordinates will be re-aligned to refine probe specificity, variant annotation, and cross-species comparability with *T. pratense*, further improving the accuracy of future mapping and genomic prediction studies.

Beyond SNP arrays, our genomic resource can also be adapted to flexible, lower-cost systems like DartTag (Diversity Arrays Technology) and Kompetitive Allele-Specific PCR (KASP) assays offered by LGC Genomics. Both approaches rely on high-quality, validated SNP catalogs for assay design. The 542K high-confidence SNP set and the ∼28K enrichment panel provide an ideal substrate for selecting subsets of markers for trait-specific genotyping. Integrating high-quality SNPs into genotyping arrays or amplicon panels is expected to greatly improve trait mapping resolution and selection accuracy. It is important to emphasize, however, that directly combining 50X and 2X coverage datasets for population genetic analyses may introduce coverage bias and skew allele frequency estimates. This was evident in our preliminary PCA analyses that showed separation of 50X and 2X samples, likely due to read depth discrepancies rather than true population structure. As such, diversity metrics, population structure inference (e.g., ADMIXTURE), and FST should rely on low-pass (2X) samples exclusively to avoid misrepresentation.

### Utility of a Mid-Density Enrichment Panel for Predictive Breeding

We evaluated a custom Twist hybrid-capture panel in our crimson clover panel of breeding materials using Khufu low-pass sequencing with reference-guided imputation. The resulting imputed (I)HapMap achieved consistently high sample- and locus-level call rates (Fig. 13), yielding a near-complete genotype matrix appropriate for breeding-scale analyses. Importantly, these metrics reflect the final imputed genotypes used in downstream inference (e.g., GRM/GP), rather than raw read depth or capture-on-target rates. The allele-frequency spectrum is consistent with predictive breeding needs: enrichment of intermediate-frequency variants (MAF ≈ 0.3-0.5) increases information content for population structure, GRM estimation, and genomic prediction, while retention of lower-frequency alleles preserves founder-haplotype diversity and supports monitoring allele-frequency trajectories across selection cycles. Relative to fixed SNP arrays, sequence-based hybrid capture is modular and updatable, whereas arrays are constrained by static marker content and can exhibit ascertainment bias driven by discovery panels and design filters that distort allele-frequency spectra and downstream LD/FST-based inference (Albrechtsen et al., 2010; Lachance & Tishkoff, 2013). Benchmarking in animal breeding further supports capture as an array-comparable platform: a TWIST capture panel in cattle produced ∼98% genotype concordance, R^2^ ≈ 0.98, relative to SNP arrays, confirming suitability for genomic selection pipelines (Ren et al., 2024). Compared with restriction-enzyme–based GBS, hybrid capture concentrates sequencing effort on predefined loci, improving locus stability and reducing dropout, an advantage for routine, cycle-to-cycle breeding operations that require consistent marker sets (Poland et al., 2012). The availability of approximately 50k informative common markers and minor allele frequency (MAF) analysis showed that 69.3% of loci were polymorphic at MAF ≥ 0.05, provides a practical foundation for breeding applications. This position the crimson clover twist panel as a practical mid-density genotyping platform bridging WGS-based SNP discovery and operational predictive breeding in an obligately outcrossing forage legume.

### Insights from low-coverage (2X) imputation of high-coverage (50X) Data: A low-cost and scalable pipeline for future generations

Recognizing the obligately outcrossing and highly heterozygous nature of crimson clover, we outline a two-tier sequencing framework designed to balance cost efficiency, genomic resolution, and cross-generational continuity within an active breeding program. This framework combines high-coverage whole-genome sequencing (50X) of selected founder accessions from the 194 lines that established the Auburn crimson clover Breeding Program in 2022-2023 (C_0_) with planned low-pass resequencing (∼2X) of subsequent breeding cycles (e.g., C_1_-C_3_). This design follows the low-coverage imputation paradigm of Pasaniuc et al. (2012), which demonstrated that accurate genotype recovery can be achieved from sparse sequencing when imputed against a dense haplotype reference panel, providing a scalable and economical alternative to conventional high-depth resequencing. The 50X founder sequences generated a high-confidence SNP discovery panel containing approximately 542,790 variants, which serves as a haplotype reference scaffold for future imputation of low-pass data.

After stringent cross-cohort harmonization (coordinate normalization, REF/ALT alignment, and exclusion of ambiguous polymorphisms), 488,078 loci were retained as candidates for imputation. Using Beagle v5.4, we evaluated imputation performance by imputing low-pass genotypes against the founder haplotypes, achieving a mean imputation accuracy of DR^2^ ≈ 0.95 (Figure 13), with downstream analyses restricted to DR^2^ ≥ 0.8 and MAF ≥ 0.01. This validation demonstrates the feasibility of applying low-pass sequencing with imputation as a routine genotyping strategy in future breeding cycles, enabling genome-wide marker coverage for GWAS, genomic prediction, and selection decisions at substantially reduced cost. Because crimson clover currently lacks a species-specific reference genome, potential mapping or representation biases were minimized through conservative site filtering and cross-species harmonization to the *T. pratense* assembly. Importantly, this framework is extensible to sub-1X coverage (∼0.8X), allowing future breeding populations to be genotyped efficiently while maintaining high accuracy for common variants (MAF ≥ 0.05). Embedding this approach within each selection cycle will enable continuous, genome-wide genotyping to inform long-term population improvement and cultivar development. The imputed call set (*2XP.imputed.ref.hier.vcf.gz*) is provided as supplementary material to support reproducibility and reuse of the reference panel.

### Reference Genome Limitations and Future Directions

A limitation of our current study is the alignment of WGR to the red clover reference genome. As a result, a portion of reads may potentially lead to the underrepresentation of variants unique to crimson clover. To address this limitation, we have initiated a de novo genome assembly for crimson clover. This reference genome will enable more accurate variant discovery, structural variant detection, and comprehensive annotation of genes and regulatory elements relevant to agronomic traits.

## CONCLUSION

This study delivers the first genome-wide genomic resource for crimson clover and establishes a practical foundation for both modern breeding and genebank-enabled crop improvement. Through WGR of a globally representative panel, we generated a high-confidence catalogue of genome-wide variation that far exceeds the resolution of previous marker-based studies. These data resolve population structure, capture rare and private alleles, and provide a comprehensive view of standing genetic diversity within the species.

From a breeding perspective, these resources position crimson clover from a historically under-resourced species to one that can fully participate in predictive and data-driven improvement pipelines. The development of a publicly available mid-density (28,913 SNPs) hybrid-capture enrichment panel bridges discovery genomics with operational breeding. The panel has been synthesized, validated, and deployed within the Auburn crimson clover breeding program, enabling direct integration of genome-wide marker information into recurrent selection pipelines. In addition to supporting genomic prediction, this platform enables tracking of allele-frequency changes across selection cycles and more effective management of genetic diversity. A key requirement for sustained genetic gain in open-pollinated, highly heterozygous species.

Equally important is the value of these resources for gene banks and germplasm management. The genome-wide SNP dataset enables robust characterization of accessions, identification of genetically distinct or underrepresented materials, and detection of redundancy within collections. The availability of standardized and transferable markers further supports interoperability between gene banks and breeding programs, facilitating more effective utilization of conserved diversity. These advances align crimson clover with genotyping platform and SNP panel resources already established in soybean, alfalfa, and peanut, bringing this species to parity with other well-resourced legumes and enabling long-term, genomics-enabled improvement.

## Supporting information

Here

## ABBREVIATIONS

WGR: whole-genome re-sequencing
LD: linkage disequilibrium
PYT: Preliminary Yield Trial
MAF: minor allele frequency
PCA: principal component analysis
SNP: single-nucleotide polymorphism
SPT: Space Plant Trial
BLUP: Best Linear Unbiased Predictor
AU: Auburn University
UF: University of Florida
PYT: Preliminary Yield Trial
DR^2^: Dosage coefficient of determination (Beagle imputation accuracy metric)
AUHPC: Auburn University High Performance Computing
JGI: Joint Genome Institute
WGS: Whole genome sequencing.

## Acknowledgments

The work (proposal: 10.46936/10.25585/60008632) conducted by the U.S. Department of Energy Joint Genome Institute (https://ror.org/04xm1d337), a DOE Office of Science User Facility, is supported by the Office of Science of the U.S. Department of Energy operated under Contract No. DE-AC02-05CH11231. Sequencing for this study was performed through the JGI Community Science Program (Proposal ID: 509088). Computational analyses were conducted using the Auburn University Easley High-Performance Computing Cluster. We thank the JGI sequencing and data management teams for their support, and Kerrie Barry et al., for coordination of data release and project guidance. We also acknowledge Hudson Alpha and TWIST Bioscience, including Sharon Reikhav and Mary Cook et al, for technical assistance with development of the crimson clover enrichment capture panel and genotyping. Portion of genotyping activity were supported by supported by the U.S. Department of Agriculture, National Institute of Food and Agriculture (USDA-NIFA), AFRI Competitive Grant 2023-67014-39449 awarded to Auburn University.

## Supplemental Material

Attached bed file format we included for enrichment panel (CrimsonClover30K_AuburnUniversity.bed), upon request is the high and low coverage and reference panel imputed SNPs (due to huge size).

## Data availability

Raw whole-genome sequencing data generated in this study have been deposited in the NCBI Sequence Read Archive (SRA) under the umbrella BioProject accession PRJNA1402467 (https://www.ncbi.nlm.nih.gov/bioproject/1402467), which links all associated data-level BioProjects corresponding to individual accessions. Processed variant datasets and resources related to the mid-density genotyping enrichment panel are provided in the Supplementary Materials. Additional processed datasets supporting the conclusions of this study are available from the corresponding author upon request.

## Author Contribution

Author Contributions: Mark Philip Castillo: Conceptualization; investigation; data curation; formal analysis; pipeline design: writing - original draft; writing - review & editing. Oluwaseye Gideon Oyebode: Conceptualization; data curation; writing - review & editing. Jayson Talag: Sequencing support. Victoria Bunting: Sequencing support. Navneet Kaur: Sequencing support. Paul Doran: Enrichment panel support and consultation; Kerrie W. Barry: Funding acquisition; data management. Jeremy Schmutz: Sequencing support; Funding acquisition; Brandon Schlautman: Funding acquisition. Alan Humphries: Funding acquisition; writing - review & editing: Kioumars Ghamkhar: Funding acquisition; writing - review & editing: Virginia M. Moore: Funding acquisition; writing - review & editing. Esteban Rios: Funding acquisition; germplasm source; writing - review & editing. Alex Harkess: Funding acquisition; sequencing support; writing - review & editing. Marnin Wolfe: Supervision; conceptualization: project administration; resources; funding acquisition; writing - review & editing.

